# Reprogramming the genome of M13 bacteriophage for all-in-one personalized cancer vaccine

**DOI:** 10.1101/2024.11.22.624916

**Authors:** Shengnan Huang, Yanpu He, Allison Madow, Huaiyao Peng, Mirielle Griffin, Jifa Qi, Mantao Huang, Heather Amoroso, Riley Abrashoff, Nimrod Heldman, Angela M. Belcher

**Affiliations:** The David H. Koch Institute for Integrative Cancer Research, Massachusetts Institute of Technology, Cambridge, MA, 02139, USA; Department of Biological Engineering, Massachusetts Institute of Technology, Cambridge, MA, 02139, USA; Department of Brain and Cognitive Sciences, Massachusetts Institute of Technology, Cambridge, MA, 02139, USA; Department of Nuclear Science and Engineering, Massachusetts Institute of Technology, Cambridge, MA, 02139, USA; Biopolymers Core Lab, The David H. Koch Institute for Integrative Cancer Research, Massachusetts Institute of Technology, Cambridge, MA, 02139, USA; Department of Materials Science and Engineering, Massachusetts Institute of Technology, Cambridge, MA, 02139, USA

## Abstract

Peptide-based vaccines face limitations in immunogenicity and stability, and challenges in co-delivering antigens and adjuvants effectively. Virus-based nanoparticles, particularly M13 bacteriophage, present a promising solution due to their genetic modifiability, intrinsic adjuvanticity, and efficient antigen presentation capabilities. Here we developed a programmable M13 phage-based personalized cancer vaccine enabling single-step antigen-adjuvant assembly. Specifically, we designed a reprogrammed (RP) phage platform that precisely regulates Toll-like receptor 9 activation by programming its genome sequence and modulates antigen density through genetic engineering. Vaccination studies with RP phages demonstrated that the immune response could be modulated by fine-tuning the adjuvanticity and antigen density, revealing an optimal antigen dose and adjuvanticity for maximum vaccine efficacy. The RP phage induced a remarkable 24-fold increase in neoantigen-specific CD8^+^ T cells and eradicated established MC-38 tumors when combined with anti-PD-1 therapy. These findings highlight the RP phage’s potential as a powerful nanovaccine platform for personalized cancer vaccines.

## Introduction

Peptide-based vaccines exhibit inherently low immunogenicity and biochemical stability^1^. Moreover, difficulties in the co-delivery of antigens and adjuvants to secondary lymphoid organs, as well as the effective uptake of the vaccine by antigen-presenting cells (APCs) remain major barriers to optimizing vaccine efficiency^2^. To address these challenges, various nanovaccine platforms, such as DNA origami, lipoprotein nanoparticle, and polymer nanoparticle, have been developed to co-assemble the antigen and adjuvant^3–8^, but approaches that could enable single-step co-assembly would be attractive. In this regard, virus-based nanoparticles are promising vaccine platforms for infectious diseases and cancer, owing to their intrinsic adjuvanticity, antigen display versatility, and ability to deliver antigens to APCs^9–12^. Among them, bacteriophages, especially the M13 bacteriophage, stand out due to their genetic manipulability, large antigen display area, favorable safety profile, high stability, and straightforward manufacturing, making them highly suitable for clinical translations.^13–18^

M13 phage is a type of filamentous bacteriophage composed of a single-stranded DNA (ssDNA) encased in capsid proteins. It is ∼ 7 nm in diameter with a length determined by the ssDNA size, which can be controlled by a phagemid^19^. M13 phage has been widely used in phage display to select proteins with high binding affinity towards the target substrates^20^. For vaccine applications, its straightforward genetic system allows the display of significant quantities of antigen peptides on the major coat protein pVIII or minor coat protein pIII^15,21,22^. Additionally, the unmethylated cytosine-phosphate-guanine (CpG) motifs within the ssDNA genome activate Toll-like receptor 9 (TLR9)^23^, contributing to its inherent adjuvanticity^24^. Current research on the M13 phage-based vaccine focuses on modifying capsid proteins for antigen display and targeting^25–29^. Few studies address systematic modulation of antigen density on the phage capsid^30,31^. While the natural adjuvanticity of phage particles has been exploited, efforts to enhance it remain limited. These factors are critical for inducing strong immune responses, especially for weak antigens that exhibit low binding affinity towards the major histocompatibility complex (MHC) molecules and/or weak peptide-MHC (pMHC)/T cell receptor (TCR) interactions^32,33^.

In this study, we aimed to develop a programmable M13 phage-based nanovaccine by modulating the adjuvanticity and antigen density (Fig. 1). We engineered a reprogrammed (RP) phage platform by programming its ssDNA sequence for precise regulation of TLR9 activation and controlled the antigen display density through genetic engineering. By modulating both adjuvanticity and antigen density of the RP phage, our results demonstrated that increased adjuvanticity significantly boosted the frequency of antigen-specific CD8^+^ T cells, revealing an optimal antigen density and adjuvanticity. Leveraging these insights, we developed an RP phage-based personalized cancer vaccine targeting neoantigens (neoAgs). The RP phage elicited a remarkable 24-fold increase in the frequency of neoAg-specific CD8^+^ T cells, significantly inhibited tumor growth, and when combined with immune checkpoint inhibitors (ICIs), eradicated established tumors in ∼ 75% of treated animals. These findings highlight the potential of the RP phage platform as a promising strategy for personalized cancer vaccines and combinatorial immunotherapies.

**Fig. 1.**
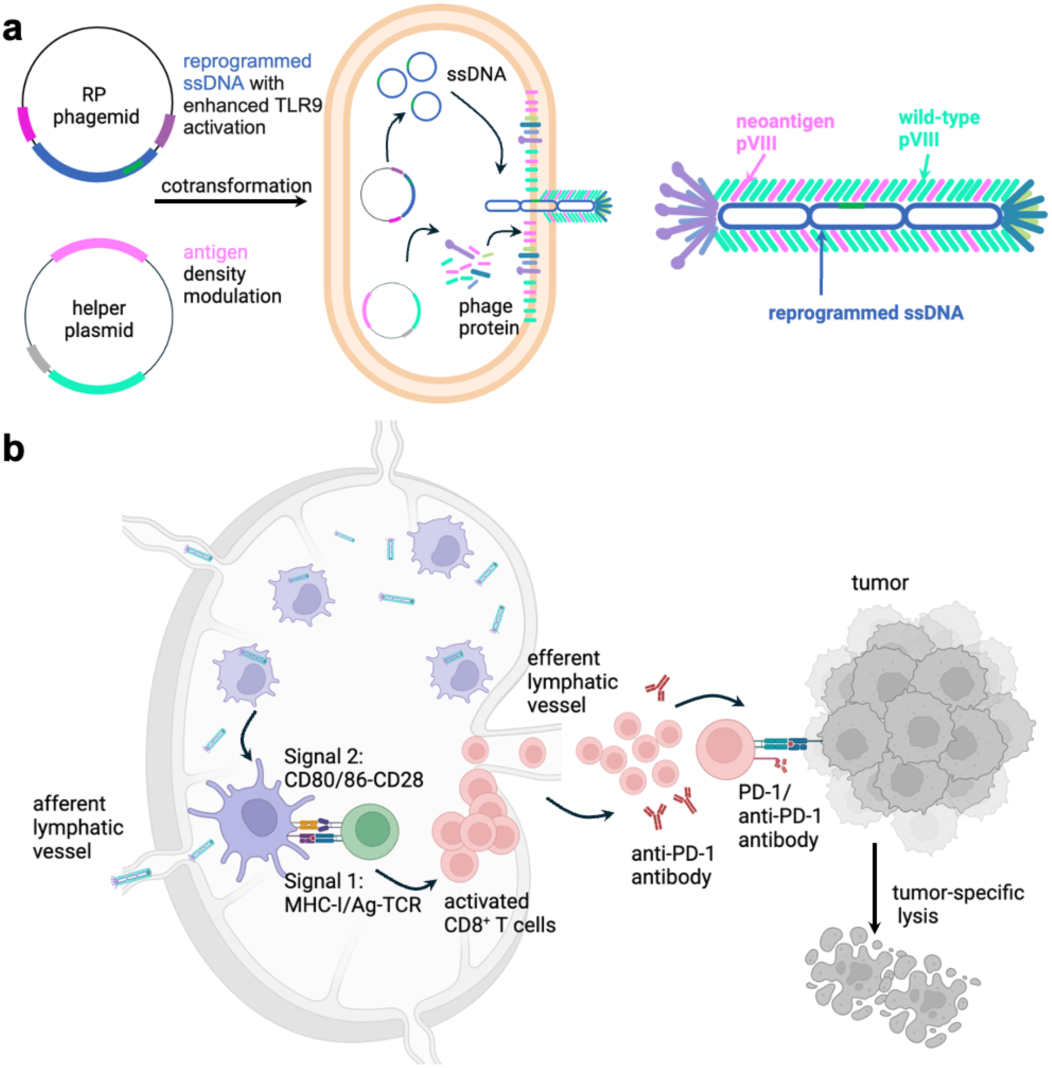
| RP phage for personalized cancer vaccine. **a**, RP phage amplification by co-transforming the RP phagemid with reprogrammed ssDNA and the helper plasmid encoding the neoantigen pVIII. The sequence of the reprogrammed ssDNA is programmed to enhance the activation of the TLR9. The helper plasmid is designed to modulate the neoantigen pVIII display on the M13 phage capsid. **b**, RP phages are efficiently taken up by APCs in the lymph node, eliciting strong antigen presentation (Signal 1) and APC maturation (Signal 2), which in turn present the antigen to the CD8^+^ T cells, resulting into the activation of antigen-specific CD8^+^ T cells. The activated antigen-specific CD8^+^ T cells recognize the cancer cells and eradicate the target cancer cells in peripheral tissue. Combining with the immune checkpoint inhibitors, such as anti-PD-1 antibodies, further amplifies the potency of the RP phage-based personalized vaccine for cancer immunotherapy.

## Results

### Modulating the adjuvanticity of the M13 phage with a custom algorithm

The phagemid-helper plasmid system was utilized to amplify M13 phages, which enabled precise and independent control over the sequence of the enclosed ssDNA (Fig. 2a)^19^. We aimed to enhance the adjuvanticity of the M13 phage by incorporating more CG dimers into the ssDNA to increase the TLR9 activation^34,35^. Specifically, we developed an algorithm (Methods section) to precisely program the ssDNA sequence. We created a series of ssDNAs of the same size (1,415 nucleotides), with varying CG fractions of 0%, 9%, and 27%, designated as CG0, CG09, and CG27, respectively. RP phages encapsulating the altered ssDNAs were amplified by combining the reprogrammed phagemids with a helper plasmid encoding the wild-type (WT) pVIII. Since the length of the M13 phage is proportional to the ssDNA size, the RP phages developed were all ∼ 200 nm in length (Fig. 2b), differing only in the ssDNA sequence. The TLR9 activation was increased by ∼ 1.7-fold (p<0.0001) from CG0 to CG27 (Fig. 2c). To further enhance TLR9 activation, we incorporated more potent canonical CpG hexamers (AACGTT and GACGTT)^36^ into the reprogrammed ssDNA. When mutating 40% of the CG dimers in CG27 into the CpG hexamers, CpG40 RP phages with a length of ∼ 220 nm were generated, which achieved ∼ 60% increase in TLR9 activation (Fig. 2d, p<0.0001) compared to the CG27 phages in an independent study, leading to an overall ∼ 2.7-fold (CG0 vs. CpG40) increase in TLR9 activation. Thus, we demonstrated that the adjuvanticity of M13 phage can be precisely modulated by programming its ssDNA sequence using our custom algorithm.

**Fig. 2.**
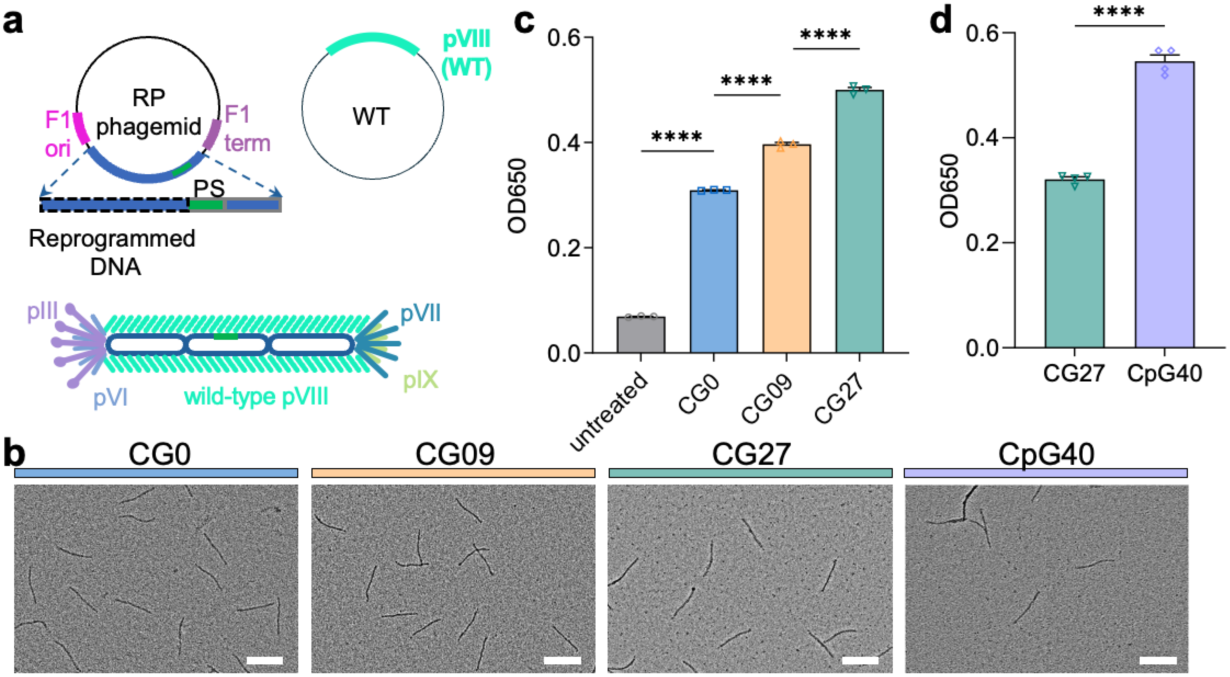
| Modulating the adjuvanticity of the M13 phage by programming its ssDNA sequence. **a**, The ssDNA sequence in between the F1-ori and F1-term (black dashed outline) of the M13 phage was reprogrammed to increase its stimulation of the TLR9 by incorporating more TLR9 agonists. **b**, Transmission electron microscopy (TEM) images of the RP phages showed the length of the RP phage was ∼ 200 nm. Scale bars represent 200 nm. **c**, The activation of TLR9 increased with increasing CG fraction in the ssDNA when studied using HEKBlue mTLR9 cell assay. **d**, CpG40 phagemid was developed when replacing 40% of the CG dimers in the CG27 phagemid with the canonical CpG hexamers. The as-assembled RP phages exhibited higher TLR9 activation. The data show mean ± s.e.m. from a representative experiment of 2-3 independent experiments. ****p<0.0001, analyzed by one-way ANOVA (**c**) with Bonferroni post hoc test, two-tailed unpaired Student’s t-test (**d**).

### Tuning the antigen pVIII density

For our initial vaccine studies, we tried to express SIINFEKL, a model antigen known to stimulate T cells through binding to the H-2K^b^ of MHC-I molecules, at the N-terminus of pVIII, which is ideal for displaying antigen peptides in large quantities. We constructed a helper plasmid, rEES, containing a recombinant pVIII (rpVIII) under the control of the Lac operon and Tac promoter (Fig. 3a, second row) to express SIINFEKL at the N terminus of rpVIII. The resulting M13 phages would be a mosaic phage consisting of WT and SIINFEKL pVIIIs, which was confirmed by matrix-assisted laser desorption/ionization time-of-flight mass spectrometry (MALDI-TOF MS, Fig. 3b, first and second rows). The SIINFEKL pVIII peak at 6142 Da was detected along with the WT pVIII peak at 5239 Da. Next, we quantified the SIINFEKL pVIII display ratio ( the proportion of SIINFEKL pVIIIs relative to the total pVIIIs) using high-performance liquid chromatography (HPLC), which is more straightforward compared to the N-terminal sequence analysis^37,38^. HPLC collection and MALDI-TOF MS analysis (Supplementary Fig. 1) confirmed that the HPLC peak at ∼ 51.3 min corresponded to the WT pVIII, while the shoulder peak following it represented the SIINFEKL pVIII (Fig. 3c). The HPLC curve was fitted using a custom Double-Gaussian fitting algorithm assuming that each HPLC peak could be modeled as a combination of two Gaussian distributions, and the SIINFEKL pVIII display ratio was quantified to be ∼ 15% (Fig. 3c, second row), which was independent of the phagemids (Fig. 3d).

**Fig. 3.**
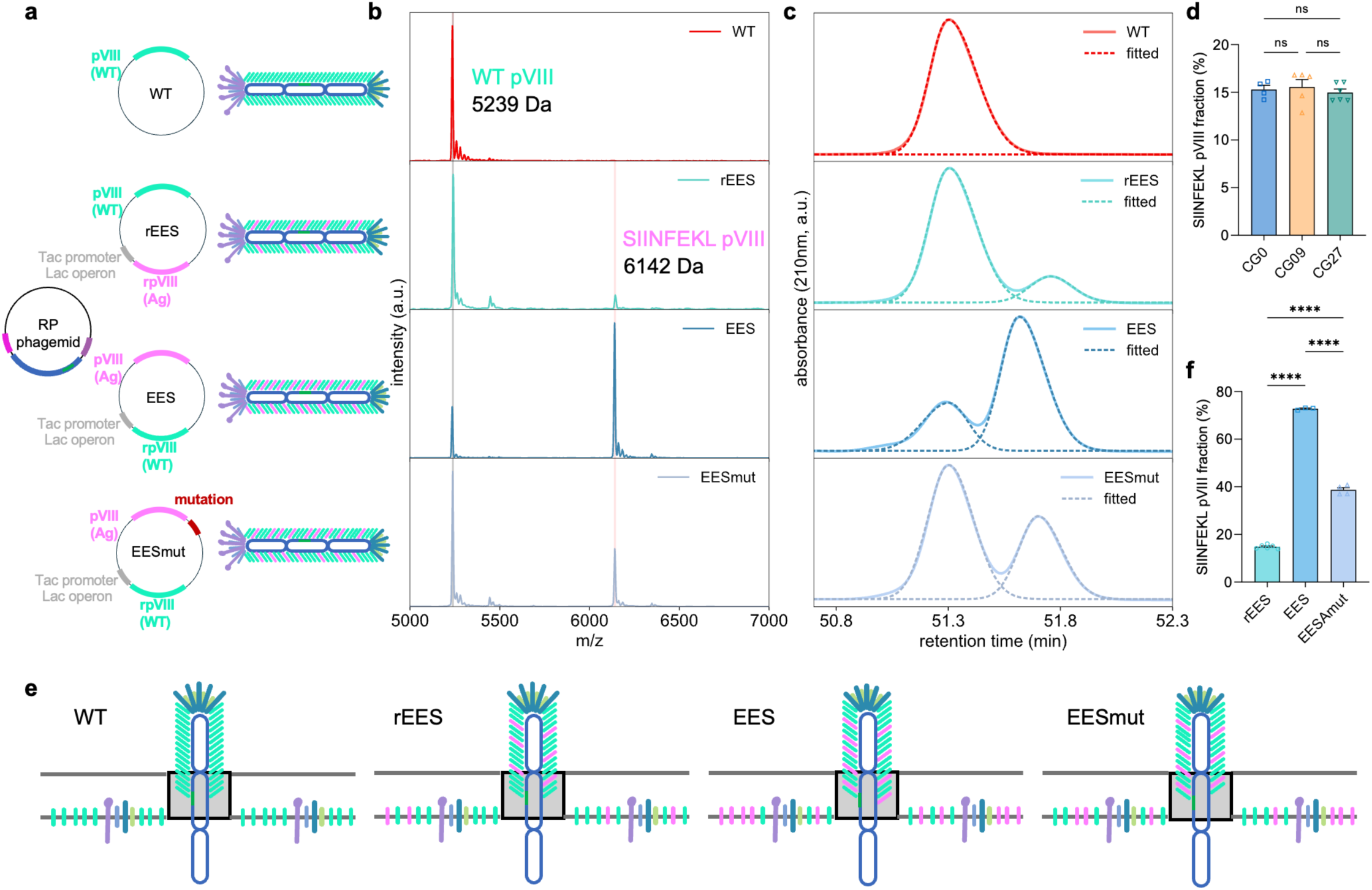
| Tuning the antigen display density on the RP phage by controlling the proportion of antigen pVIII relative to wild-type pVIII. **a**, SIINFEKL pVIII-displaying RP phages were amplified by combining the RP phagemid and the SIINFEKL pVIII-expressing helper plasmid: rEES, EES and EESmut. **b**, Confirmation of the SIINFEKL pVIII display on the RP phage by MALDI-TOF mass spectrometry. **c**, Quantification of the SIINFEKL pVIII display ratio on the RP phage by HPLC. **d**, SIINFEKL pVIII display ratio of RP phages amplified with various RP phagemids and the rEES helper plasmid was independent of the phagemid. **e**, Controlling the SIINFEKL pVIII display density on the RP phage. **f**, A summary of the SIINFEKL pVIII display ratios of RP phages amplified with various helper plasmids. The data show mean ± s.e.m.. ns: none significance, ****p<0.0001, analyzed by one-way ANOVA (**d**, **f**) with Bonferroni post hoc test.

Then we aimed to increase the antigen display ratio, hypothesizing that higher antigen density could increase pMHC density and subsequent antigen presentation to TCRs^32,39^. During M13 phage assembly, all the capsid proteins are synthesized in the *E. coli* cytoplasm and transported to the inner membrane for assembly (Fig. 3e)^40^. We hypothesized that increasing the proportion of antigen pVIII relative to the WT pVIII would increase its assembling probability, thereby increasing the display ratio (Fig. 3e). To achieve this, we swapped the expression sites of WT and antigen pVIII in a new helper plasmid EES (Fig. 3a, third row). MALDI-TOF MS (Fig. 3b, third row) and HPLC results (Fig. 3d, third row) showed an increased display ratio of ∼ 73% (Fig. 3f), however, the incorporation of antigen pVIIIs disrupted phage assembly and reduced amplification yield. We then sought to identify any favorable mutations in the helper plasmid EES to increase the amplification yield without significantly compromising the SIINFEKL pVIII display ratio. By continuously monitoring the amplification yield and display ratio, a mutation preceding the start codon of the SIINFEKL pVIII was identified, resulting in the helper plasmid EESmut (Fig. 3a, bottom row), which yielded an SIINFEKL pVIII display ratio of ∼ 39% (Figs 3b,c,f), while significantly enhancing the phage yield. To the best of our knowledge, this mutation, potentially a ribosome binding site mutation based on its location, has not been previously reported. There is a tradeoff between the peptide length and its pVIII N-terminus display ratio on the M13 phage^41^. For a peptide of 10 amino acid residues (EESIINFEKL), a display ratio of ∼ 39% is considered high. Thus, we have developed a novel approach to control the antigen density, potentially enabling fine-tuned modulation of the immune response based on the antigen dose.

### RP phages accumulated in lymph nodes and captured by APCs

As antigen presentation and T cell priming orchestrate in lymph nodes (LNs)^2^, we next characterized the RP phage (∼ 200 nm long) drainage into the draining LNs (dLNs) by conjugating it with fluorescein (Supplementary Fig. 2). 24 hours after the subcutaneous injection, RP phages showed significantly greater accumulation in inguinal (IN) LNs than the free antigen and adjuvant combinations (free SIINFEKL and CpG, freeS/CpG, Fig. 4a, p<0.0001), while freeS/CpG showed minimal LN drainage. RP phages also tended to accumulate in the axillary (AX) LNs (∼ 20% of IN LNs) with minimal accumulation in the brachial (BR) LNs. Similarly, histological sections of the dLNs revealed minimal detectable free antigen, while RP phages accumulated in the subcapsular sinus and interfollicular areas (Fig. 4b). The enhanced LN trafficking of RP phages could be attributed to their nanoscale dimensions, which exploit the physiological characteristics of lymphatic drainage^42^. Within the immune cell populations of the dLNs, RP phages were mostly associated with macrophages (Mφs, CD3^-^/B220^-^/CD11b^+^/CD11c^-^/F4/80^+^, ∼ 23.1%), dendritic cells (DCs, CD3^-^/B220^-^/CD11b^-^/CD11c^+^, ∼ 18.1%) and B cells (CD3^-^/B220^+^, ∼ 3.1%) as in Fig. 4c, all of which are professional APCs (FACS gating approach in Supplementary Fig. 3), consistent with Refs. ^2,43^. Importantly, RP phages led to the activation of Mφs and DCs, as indicated by the elevated expression of co-stimulatory molecules CD80 and CD86 (Fig. 4d). Compared to the freeS/CpG group, the RP phage group exhibited a ∼ 4-fold increase in CD80^+^CD86^+^ DCs (p<0.0001) and a ∼ 2-fold increase in CD80^+^CD86^+^ Mφs (p<0.0001), while freeS/CpG had minimal effect on Mφ and DC maturation compared to 1× PBS. Overall, the results suggested that RP phages efficiently drained to the dLNs, preferentially accumulated in professional APCs, and substantially enhanced their activation, thereby providing the foundation for effective antigen presentation and subsequent T-cell activation in the dLNs.

**Fig. 4.**
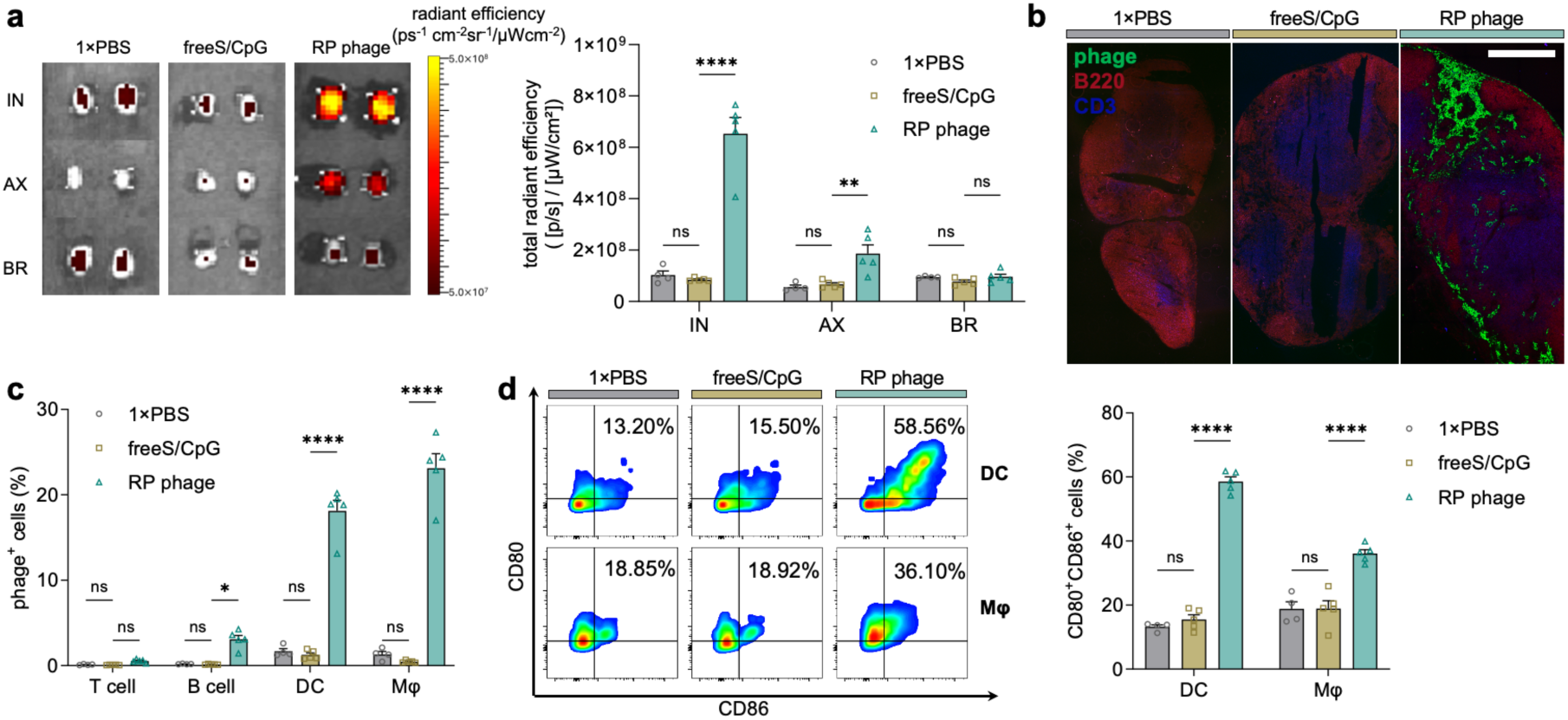
| Accumulation of RP phages in the draining lymph nodes and professional APCs. **a**, RP (CG27/rEES) phages preferentially accumulated in the inguinal lymph node compared to free antigen as studied by the in vivo imaging system (IVIS). IN: inguinal, AX: axillary, BR: brachial**. b**, Spatial distribution of RP phages in draining lymph nodes. Scale bar represents 500 μm. **c**, RP phages were mostly associated with macrophages (Mφs) and dendritic cells (DCs). **d**, Activation of Mφs and DCs by RP phages as indicated by the upregulation of CD80 and CD86. The data labeled is the mean of each group. The data show mean ± s.e.m. from 2-3 independent experiments (n= 4-5). ns: none significance, *p<0.05, **p<0.01, and ****p<0.0001, analyzed by two-way ANOVA (**a**, **c**, **d)** with Bonferroni post hoc test.

### Robust and durable CD8^+^ T cell response elicited by RP phages

We then modulated the vaccine efficacy by controlling the adjuvanticity and/or antigen density of the RP phages. Female C57BL/6 mice were immunized with RP phages of varying adjuvanticity (CG0, CG09, CG27, Fig. 2c) but with the same ssDNA size (Fig. 2b) and antigen density (rEES, Fig. 3d) (Fig. 5a). The results showed a marked increase in the frequency of SIINFEKL-specific CD8^+^ T cells with increasing adjuvanticity, from ∼ 10.5% to ∼ 17.4% (CG0 vs CG27, p<0.05, Fig. 5b). Upon challenge with 0.3M B16F10-OVA cells, mice vaccinated with RP phages (CG27/rEES) showed slower tumor growth and extended survival compared to other groups (Figs 5c, d). However, further increasing the RP phage adjuvanticity (CG27/rEES vs CpG40/rEES) without changing the antigen density did not boost the vaccine efficacy (Fig. 5e). Moreover, increasing both the antigen density (CG27/rEES vs CG27/EESmut, Fig. 5f) and adjuvanticity (CG27/EESmut vs. CpG40/EESmut, Fig. 5f) failed to enhance the vaccine efficacy. Collectively, these results suggest that there is an optimal dose for antigen and adjuvant to elicit the maximum immune response from nanovaccines and that exceeding these levels may not further enhance efficacy. Additionally, free S/CpG induced minimal SIINFEKL-specific CD8^+^ T cell response and provided no protection against tumor invasion when challenged with 0.1 million B16F10-OVA cells (Figs 5g,h). In contrast, 5 out of 6 mice in the RP phage groups remained tumor-free 40 days post-inoculation (Fig. 5i).

**Fig. 5.**
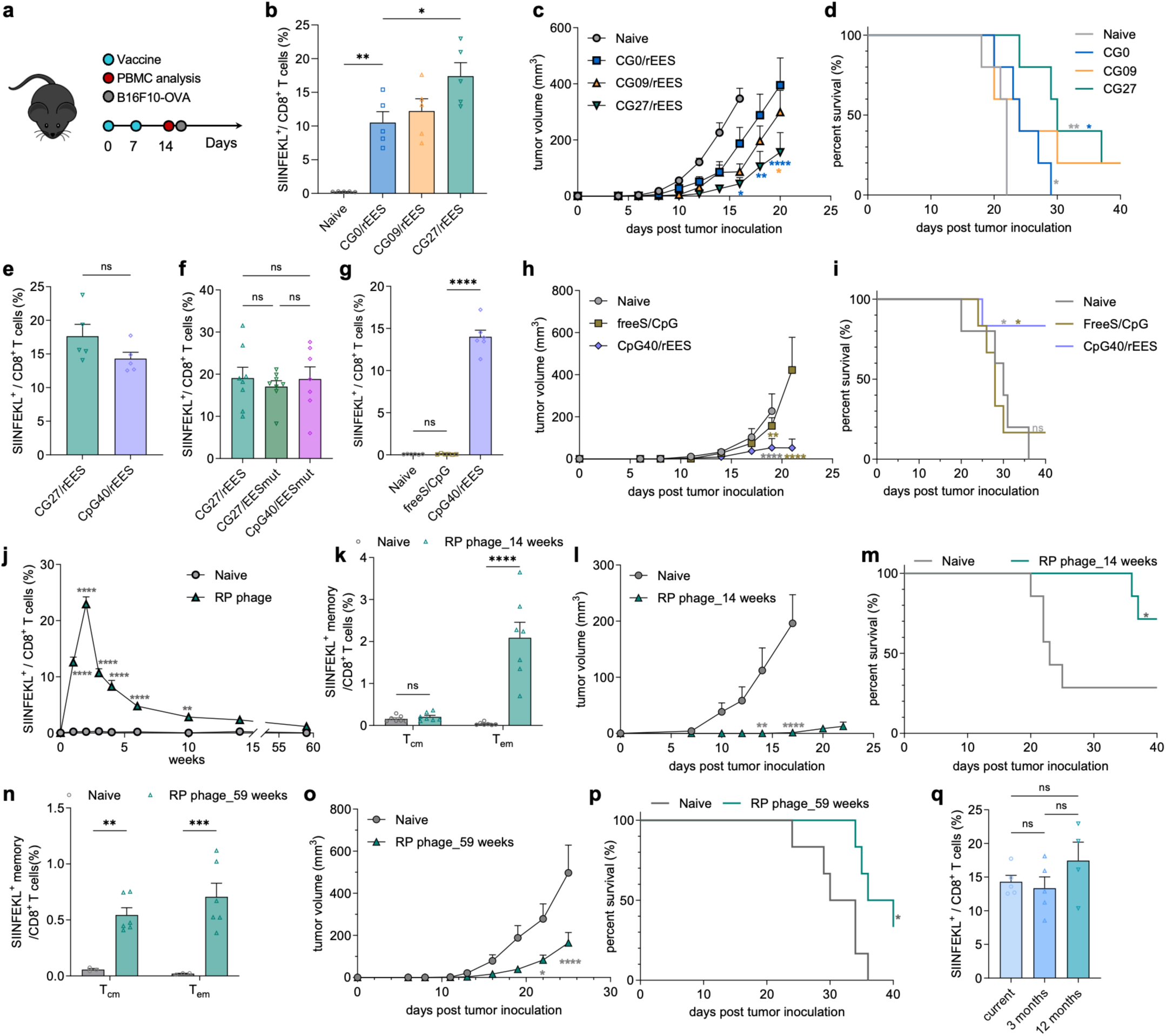
| Robust CD8^+^ T-cell response and long-lasting memory elicited by RP phages. **a**, Vaccination dosing schedule of the RP phages with C57BL/6 mice. Two doses of RP phage were administered via tail base injection on day 0 and day 7. Peripheral blood was analyzed on day 14 for SIINFEKL-specific CD8^+^ T cells and B16F10-OVA cells were inoculated on day 15. **b**, The frequency of SIINFEKL-specific CD8^+^ T cells in the peripheral blood increased with increasing adjuvanticity of the RP phages amplified by the same helper plasmid (rEES) and various phagemids (CG0/09/27). **c**, On day 15, mice were challenged with 0.3M B16F10-OVA cells subcutaneously on the left flank, and tumor growth was monitored over time. **d**, Survival study of the mice in (**c**) showed that RP phage vaccination substantially prolonged the survival. **e**, When increasing the adjuvanticity of the RP phages (CG27 vs CpG40), SIINFEKL-specific CD8^+^ T cell response did not increase. **f**, Increasing antigen display density (rEES vs EESmut) and adjuvanticity (CG27 vs CpG40) did not enhance the SIINFEKL-specific CD8^+^ T cell response. **g**,**h**,**i**, RP phage demonstrated better vaccination efficacy compared to free antigen and adjuvant. **g**, Free SIINFEKL peptide and CpG formulation (freeS/CpG) elicited minimal T-cell response. RP phages significantly slowed down the tumor progression (**h**) and prevented tumors from developing (**i**) compared to freeS/CpG. **j**, Fraction of SIINFEKL-specific CD8^+^ T cells over time post RP phage administration on day 0 and day 7. **k**,**l**,**m**, SIINFEKL-specific CD8^+^ T cell memory study at week 14. **k**, Fraction of SIINFEKL-specific central memory CD8^+^ T cells (T_cm_) and effector memory CD8^+^ T cells (T_em_) at week 14. Vaccination with RP phages slowed down tumor progression (**l**) and prevented tumors from developing (**m**) for mice at week 14. **n**,**o**,**p** SIINFEKL-specific CD8^+^ T cell memory study at week 59. **n**, Fraction of T_cm_ and T_em_ at week 59 in an independent study. Vaccination with RP phages slowed down tumor progression (**o**) and extended the survival (**p**) of mice at week 59. **q**, RP phage showed comparable vaccination efficiency after 1 year of storage at 4 °C. The data show mean ± s.e.m.. ns: none significance, *p<0.05, **p<0.01, ***p<0.001 and ****p<0.0001, analyzed by one-way ANOVA (**b**,**f**,**g**,**q**) or two-way ANOVA(**c**,**h**,**j**,**k**,**l**,**n,o**) with Bonferroni post hoc test and long-rank test (**d,i**,**m**,**p**).

Next, we examined the durability of the antigen-specific CD8^+^ T cells elicited by RP phages. The frequency of the SIINFEKL-specific CD8^+^ T cells first expanded before contracting, peaking one week after the second dose (Fig. 5j). By week 14, the frequency reached ∼ 2.3%, decreasing to ∼ 1.2% by week 59 in an independent study. The temporal response of the antigen-specific CD8^+^ T cells elicited by RP phages resembles a typical T cell response pattern post antigen stimulation with expansion, contraction, and memory phases^44^. At week 14, phenotype analysis of the SIINFEKL-specific CD8^+^ T cells showed that ∼ 90% (2.1% out of 2.3%) were effector memory T cells (T_em_, CD8^+^/SIINFEKL^+^/CD44^+^/CD62L^-^) with negligible central memory T cells (T_cm_, CD8^+^/SIINFEKL^+^/CD44^+^/CD62L^+^) (Fig. 5k, FACS gating strategy in Supplementary Fig. 4). Upon challenge with 80k B16F10-OVA cells, RP phage significantly slowed the tumor progression compared to age-matched naïve mice, with 5 out of 7 mice remaining tumor-free on day 40 (Figs 5l, m). By week 59, phenotype analysis of the SIINFEKL-specific CD8^+^ T cells revealed a significant shift with ∼ 42% T_cm_ (0.5% out of 1.2%) and ∼ 58% T_em_ (0.7% out of 1.2%) (Fig. 5n), indicating durable CD8^+^ T cell memory and phenotypic evolution over time^45^. Upon inoculation with 30k B16F10-OVA cells, the long-lasting SIINFEKL-specific memory CD8^+^ T cells effectively delayed tumor progression (Figs 5o,p). Moreover, RP phages stored at 4 °C for one year elicited comparable antigen-specific CD8^+^ T cell response (Fig. 5q), underscoring the exceptional stability of RP phages, which is critical for the broad vaccine distribution.

### RP phage-based personalized cancer vaccine by enhancing the adjuvanticity and antigen density

To demonstrate the versatility of our RP phage platform for personalized cancer vaccines targeting neoAgs, we engineered an RP phage displaying the Adpgk neoAg in the MC-38 colon carcinoma model. The Adpgk neoAg harbors a single epitope mutation (ASMTN<u>R</u>ELM → ASMTN<u>M</u>ELM) and is known to be presented in the MHC-I H-2D^b^ molecules^5,6^. The presence of the Adpgk mutation in the MC-38 cell was confirmed with cDNA sequencing (Supplementary Fig. 5). Utilizing the antigen pVIII density modulation approach developed in Fig. 3, we tuned the Adpgk pVIII density on the RP phages with two helper plasmids, rAE and AE (Fig. 6a). Specifically, MALDI-TOF MS verified the presence of the Adpgk pVIII peak at 6005 Da, with a notable increase when using the AE helper plasmid (Fig. 6b). HPLC (Fig. 6c, Supplementary Fig. 6) and the Double-Gaussian fitting method estimated the Adpgk pVIII display ratios at ∼ 16% and ∼ 60% for rAE and AE plasmids, respectively (Fig. 6d), which was also independent of the phagemids (Fig. 6e).

**Fig. 6.**
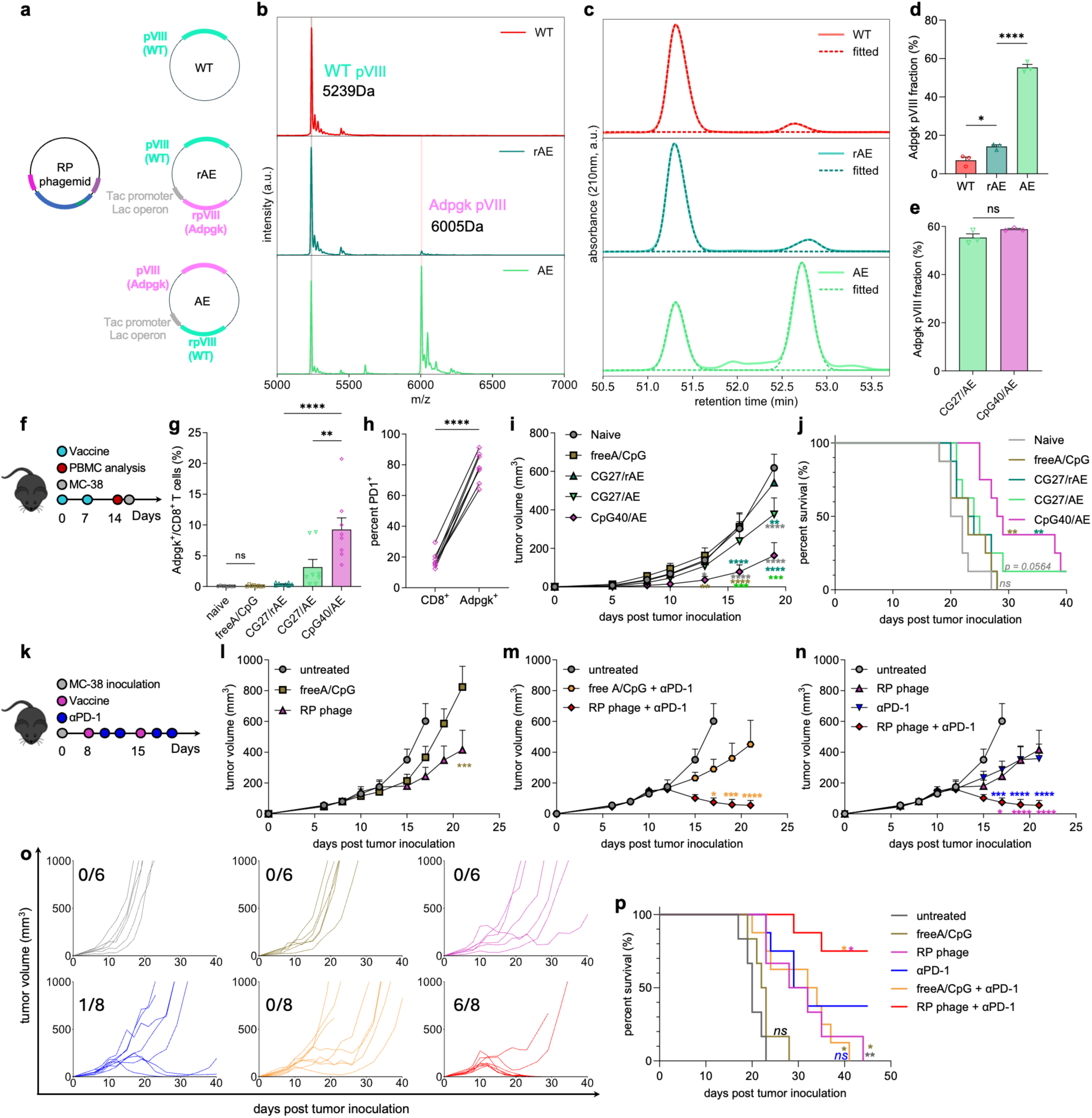
| RP phage-based personalized cancer vaccine for prophylactic and therapeutic applications. **a**, Adpgk pVIII-displaying RP phages amplified with the RP phagemid and the helper plasmids: rAE and AE**. b-e**, Quantification of the Adpgk pVIII display ratio. Confirmation and quantification of the Adpgk pVIII with MALDI-TOF **(b)** and HPLC **(c)**, respectively. **d**, Adpgk pVIII display ratio for the rAE and AE helper plasmids. **e**, The Adpgk pVIII display density was independent of the phagemids. **f-j**, Enhancing RP phage vaccination efficacy by increasing the adjuvanticity and antigen density of RP phages. **f**, Dosing schedule of the vaccination study. **g**, Increasing both the Adpgk pVIII display and adjuvanticity of the RP phage led to stronger Adpgk-specific CD8^+^ T cell response. **h**, Upregulation of the PD-1 expression in the Adpgk-specific CD8^+^ T cells. Vaccination with RP phage of CpG40/AE slowed down the MC-38 tumor growth (**i**) and prolonged the survival (**j**) of the mice. **k**-**p**, Strong synergy between RP phage and anti-PD-1 therapy led to complete tumor regression. **k**, C57BL/6 mice were subcutaneously inoculated with 0.25M MC-38 cells on day 0. For vaccine only treatment, mice were vaccinated with the RP phage or soluble vaccine via tail base injection on day 8 and 15. For anti-PD-1 antibody treatment only, mice were administered with anti-PD-1 antibodies intraperitoneally on day 10, 12, 17, and 19. For combined therapy, on day 2 and 4 after each vaccination, mice were administrated intraperitoneally with anti-PD-1 antibodies. Average tumor volume of the vaccine only treatment study (**l**), combined therapy study (**m**), and synergy study between RP phages and anti-PD-1 antibodies (**n**). **o**, Individual MC-38 tumor growth curves and (**p**) survival curves of the different study groups in (**l**, **m**, **n**). The data show mean ± s.e.m. from 2-3 independent experiments (n = 6-8). ns: none significance, *p<0.05, **p<0.01, ***p<0.001 and ****p<0.0001, analyzed by two-tailed unpaired Student’s t-test (**e,h**), one-way ANOVA (**d**,**g**) or two-way ANOVA (**i**,**l**,**m,n**) with Bonferroni post hoc test and long-rank test (**j**,**p**).

In a preliminary vaccination trial following the schedule in Fig. 6f, RP phages of varying adjuvanticity but the same antigen density (CG0/rAE, CG09/rAE, CG27/rAE, and CpG40/rAE) elicited low Adpgk-specific CD8^+^ T cell frequencies (Supplementary Fig. 7), suggesting that antigen density might be the limiting factor. To address this, we used the AE plasmid to increase the Adpgk pVIII display ratio. In vivo vaccination results demonstrated that the RP phage CpG40/AE exhibited a significant increase in Adpgk-specific CD8^+^ T cell fraction of ∼ 9.3%, representing a ∼ 24-fold enhancement compared to the CG27/rAE (∼ 0.38%, p<0.0001, Fig. 6g). Furthermore, the upregulation of the PD-1 receptor on the Adpgk-specific CD8^+^ T cells, compared to the total CD8^+^ T cells, indicated the potent stimulation of the CD8^+^ T cells by the CpG40/AE phage (Fig. 6h). When challenged with 85k MC-38 cells on day 15, CpG40/AE significantly delayed the tumor progression (Fig. 6i) and improved survival compared to the other study groups (Fig. 6j). Specifically, CpG40/AE prolonged survival by ∼ 40% relative to the naïve group (median survival 29.5 days vs 21 days, p<0.001). In contrast, vaccination with free A/CpG failed to elicit Adpgk-specific CD8^+^ T cells and inhibit the tumor progression compared to the naïve group. Thus, we developed an RP phage-based personalized cancer vaccine that demonstrated strong prophylactic efficacy by enhancing the neoAg pVIII density and adjuvanticity of the phage.

Lastly, we evaluated the therapeutic efficacy of the RP phage-based personalized cancer vaccination. Following the treatment schedule in Fig. 6k, the RP phage (CpG40/AE) significantly inhibited the tumor growth, compared to other study groups (Fig. 6l) with a median survival of 30 days, representing an increase of ∼ 50% and ∼ 33% than the untreated group (20 days) and soluble vaccine group (22.5 days), respectively. However, RP phage treatment did not result in complete tumor regression, likely due to the immunosuppression within the tumor microenvironment^5^. In order to block the immune-suppressive PD-1/PD-L1 pathway, we combined the RP phage vaccine with anti-PD-1 (αPD-1) antibodies^46^. We found that the combination therapy efficacy depended critically on the treatment timing (Supplementary Fig. 8), highlighting the importance of early tumor detection for achieving better treatment outcomes^47^. When initiating the treatment on day 8 with an average tumor size of ∼ 80mm^3^, the RP phage/αPD-1 combination significantly enhanced treatment outcomes (Fig. 6m) and demonstrated strong synergy (Fig. 6n), achieving complete tumor regression in ∼ 75% of mice (6 out of 8, Figs 6o,p). Thus, RP phage-based personalized cancer vaccine shows great potential for therapeutic treatment of cancer, particularly when combined with other treatment modalities.

## Conclusion

The study successfully demonstrates the potential of the M13 phage as a versatile and effective nanovaccine platform by optimizing both its adjuvanticity and antigen density. The novel algorithm developed for reprogramming the genome and the innovative approach to tune the antigen density have provided valuable insights into vaccine design. Using SIINFEKL as a model antigen, we showed that enhancing the adjuvanticity of the M13 phage could significantly boost the immune response, however, there was an optimum dose for both antigen and adjuvant for maximum vaccine efficiency. Leveraging these insights in developing a personalized cancer vaccine targeting the MC-38 neoAg Adpgk highlights the practical benefits of this approach, leading to drastically enhanced T-cell response and significant tumor suppression. Combining the RP phage vaccine with ICIs offers a promising strategy for more effective cancer immunotherapy. These advancements pave the way for further research and optimization in personalized cancer vaccine development.

## Acknowledgement

This work was supported by the Koch Institute Frontier Research Program, the Marble Center for Cancer Nanomedicine, and the U.S. Defense Advanced Research Projects Agency. This work was also supported in part by the Koch Institute Support (core) Grant P30-CA014051 from the National Cancer Institute. Fig. 1 was created in part by the use of BioRender.com. H.P. is a recipient of the Takada fellowship.

## Author contributions

S.H. and A.M.B conceived the idea and designed the experiments. S.H. constructed the plasmids with help from Y.H. and N.H.. S.H. and Y.H. contributed to the tetramer staining assays. S.H., A.M. and M.G. purified the phages and maintained the cell culture. S.H., A.M., and H.P. conducted the lymph node draining experiments. H.P. carried out the immunohistology staining and fluorescence microscopy. M.H. and S.H. developed the algorithms for programming the ssDNA and fitting the HPLC peaks. J.Q. contributed to the TEM images. H.A. and R.A. performed the HPLC and MALDI-TOF MS. S.H. and A.M.B analyzed the results and wrote the manuscript. All the authors reviewed the manuscript.

## Methods

### ssDNA programming algorithm and RP phagemid

The ssDNA sequences of interest were generated using a custom-developed algorithm based on a set of design principles. First, the flanking DNA sequence between the CG dimers was constrained to include the nucleotides ATCG, with the condition that it cannot start with C or end with G. Second, any repeating sequences of 9 bases or longer were addressed by mutating the A/T in the first instance of the repeating sequence to T/A, continuing this process until all repetitions were eliminated. Third, hairpin structures—defined as complementary DNA sequences longer than 12 bases—were removed by mutating A to T or T to A within these regions. Subsequently, gblocks of the designed DNA sequences were synthesized by Integrated DNA Technologies.

To construct the RP phagemid, an inho phagemid was utilized^19^. The RP phagemid features a colE1 origin of replication and confers ampicillin resistance. It includes the f1 origin and terminator, along with a packaging signal, facilitating the production of high-purity ssDNA segments with precise lengths. Specifically, the inho-phagemid was linearized at the f1 origin and prior to the packaging signal using site-directed mutagenesis, after which the gblock of the designed DNA was inserted through Gibson assembly. The specific ssDNA sequences for RP phage CG0, CG09, CG27, and CpG40 can be found in Supplementary Table 1.

### Helper plasmid

The capsid plasmid features a p15a origin of replication and confers kanamycin resistance. It was derived from the RM13f1 helper plasmid of the inho-phage system. To enable the display of the antigen of interest, a recombinant form of pVIII, regulated by the Tac promoter and the Lac operon, was inserted between the M13 gene IV and the p15a origin of replication using site directed mutagenesis and Gibson assembly. The antigen sequence was incorporated at the N-terminus of pVIII, either at the recombinant pVIII site or the wild-type pVIII site, through site-directed mutagenesis. The full pVIII sequences of the antigen pVIII-expressing helper plasmids can be found in Supplementary Table 2.

### Phage amplification

Chemically competent XL1 blue cells (200249, Agilent) were co-transformed with the helper plasmid and the RP phagemid, then plated on agar containing kanamycin (50 µg/mL) and ampicillin (100 µg/mL). A single colony was subsequently selected and cultured in 800 mL of LB broth supplemented with kanamycin (50 µg/mL) and ampicillin (100 µg/mL) for two consecutive overnight incubations at 37 °C with continuous shaking at 225 rpm. The bacterial cultures were then centrifuged at 8000 rpm (Beckman Coulter JLA 8.1000) for 1 hour to pellet the bacterial cells. The supernatants containing phage were collected, and polyethylene glycol (PEG)/NaCl solution (final concentrations: 10% w/v PEG8000 and 0.5 M NaCl) was added to promote phage precipitation at 4 °C overnight. Following precipitation, the phage supernatants were centrifuged at 8000 rpm (Beckman Coulter JLA 8.1000) for 1 hour to pellet the phage particles. The resulting phage pellets were resuspended in a solution of 1× TBS/MgCl₂/DNase I (containing 5 µg/mL DNase I, 10 mM Tris-HCl, and 10 mM MgCl₂) and incubated for a minimum of 40 minutes at room temperature on a benchtop shaker to digest any DNA contaminants. After digestion, phage precipitation was facilitated by the addition of PEG/NaCl (10 w/v% PEG8000 and 0.5 M NaCl), and the mixture was incubated on ice for at least 2 hours. The phage particles were then collected by centrifugation at 14,000 rpm for 10 minutes, and the phage pellets were resuspended in 1× PBS. Endotoxins were removed through a series of at least three washes with Triton 114. Specifically, Triton 114 (1 v/v%) was added to the phage solution, and the mixture was stirred at 4 °C for at least 1 hour, followed by centrifugation at 14,000 rpm at 37 °C for a minimum of 30 minutes to remove the Triton 114. After centrifugation, the Triton 114 precipitated out of the phage solution, and the supernatant was carefully collected without disturbing the Triton 114 layer at the bottom. Following the final centrifugation, any remaining Triton 114 was removed using SM-2 bio-beads (1528920, Bio-Rad Laboratories). The phage was then purified via CsCl gradient ultracentrifugation (1.2–1.6 g/mL gradient, SW32, 30,000 rpm, 4 °C overnight). After ultracentrifugation, the phage band, located at approximately 1.3 g/mL, was isolated and desalted through dialysis using a 10-12 kDa cut-off membrane against Milli-Q water for at least 24 hours, with frequent buffer exchanges. Finally, the phage was concentrated again with PEG/NaCl (10 w/v% PEG and 0.5 M NaCl), centrifuged, and redispersed in 1× PBS for subsequent in vitro or in vivo applications.

### High performance liquid chromatography

For antigen display density quantification with high performance liquid chromatography (HPLC), M13 phages were lysed with hot water (∼ 96°C) assisted with sodium dodecyl sulfate (2 w/v%). HPLC was performed using an Agilent Model 1100 HPLC system equipped with an Agilent G1315A DAD Detector. Most of the chromatographic separation was achieved on a YMC-Triart Bio C4 silica column (4.6mm*×*150mm, 5 µm particle size, 300Å pore size) maintained at room temperature. The mobile phase consisted of 0.05% Trifluoroacetic Acid as Buffer A and 0.043% Trifluoroacetic acid, 80% Acetonitrile as Buffer B, delivered at a flow rate of 1.5 mL/min. A gradient elution program was applied as follows: 0-5 minutes isocratic @ 5% Buffer B, 5-75 minutes linear increase to 100% Buffer B. The injection volume was 95 µL (10 µL M13 phage + 90 µL 0.1% Trifluoroacetic acid, 5 µL dead volume).

### Matrix-assisted laser desorption/ionization time-of-flight mass spectrometry

Matrix-assisted laser desorption/ionization time-of-flight mass spectrometry (MALDI-TOF MS) analyses were performed on a Bruker Microflex LRF with a N_2_ laser (337 nm) in positive ion mode. The mass spectrometer was operated in a linear mode with a mass range of 3000–8000 m/z and was calibrated to a 5729 Da standard. The laser energy was set at 60% of the maximum, and 400 shots per spectrum were averaged to improve the signal-to-noise ratio. The matrix used was sinapinic acid, prepared as a 20 mg/mL solution in 70% acetonitrile with 0.1% trifluoroacetic acid. The matrix solution was mixed with the sample in a 1:1 ratio, and 1 µL of this mixture was spotted onto the MALDI target plate using the dried droplet method. The target plate was allowed to dry at room temperature.

### In vitro TLR9 cell assay

HEKblue™ mTLR9 (InvivoGen) reporter cells were employed to investigate TLR9 activation by the RP phages. Specifically, 4×10^4^ mTLR9 cells were seeded in a 96-well plate overnight in 100 μL DMEM supplemented with 10% fetal bovine serum (FBS), 100 U/mL penicillin, and 100 U/mL streptomycin. 2 hours prior to adding phage particles, the supernatant was replaced with DMEM medium supplemented with 10% heated-inactivated FBS, 100 U/mL penicillin, and 100 U/mL streptomycin. RP phages (∼ 7.5*×*10^10^ phage particles calculated based on a ssDNA length of 7234 nucleotides) were then added to each well. To facilitate the delivery of the negatively charged ssDNA to the mTLR9 cells, the phage particles were heated-inactivated by immersing them in water at 96 °C for 10 minutes and subsequently mixed with TransIT-X2 (0.2 µL/well, Mirus Bio). Following overnight incubation, 90 µL of the supernatant was collected from each well and mixed with 90 µL of QUANTI-Blue solution (rep-qbs2, InvivoGen). The mixture was incubated at 37 °C for 15–30 minutes, and the absorbance at 650 nm was measured using a plate reader.

### Mice

All animal experiment procedures and protocols were pre-approved by the Division of Comparative Medicine (DCM) and the Committee on Animal Care (CAC) under protocol #0222-017-25 at the Massachusetts Institute of Technology. These procedures complied with the Principles of Laboratory Animal Care established by the National Institutes of Health (NIH), United States.

All the mice used in the study were 8-10 weeks-old female C57/BL6J mice purchased from the Jackson Laboratory.

### Cell lines and cell cultures

B16F10-OVA cells (provided by the Irvine Lab, Koch Institute) and MC-38 cells (obtained from the Koch Institute Cell Repository) were cultured in DMEM supplemented with 10% fetal bovine serum (FBS), 100 U/mL penicillin, and 100 U/mL streptomycin, and maintained at 37 °C in a 5% CO_2_ atmosphere.

### Lymph node draining and intra-cellular distribution studies

Fluorescein (FAM)-NHS ester (55120, Lumiprobe) was dissolved in anhydrous DMSO and reacted with the phage in 1*×*PBS overnight in the dark at room temperature. Following this reaction, excess FAM molecules were removed via dialysis (10-12 kDa cut-off membrane) against 1× PBS for at least 24 hours with frequent buffer exchange. The absorption spectrum of the resulting phage-FAM complex was measured using a DU800 (Beckman) spectrophotometer.

For the lymph node drainage study, equivalent amounts of SIINFEKL peptide conjugated with FAM (SIINFEKL-FAM, AS-64231, Anaspec) and the phage-FAM complex were prepared in 100 µL 1× PBS. Additionally, 100 µL 1× PBS was included as a control. These solutions were administered at the tail base of C57/BL6 mice. After 24 hours, the inguinal, brachial, and axillary lymph nodes were excised intact, and their fluorescence (excitation: 465 nm; emission: 520 nm) was measured using an in vivo imaging system (IVIS).

Then the draining lymph nodes were dissociated into single cells and proceeded to antibody staining with the following makers: live/dead staining (L34976, Thermofisher), CD3e (BV785, clone 145-2C11, 417-0031-82, eBioscience), B220/CD45R (PE, clone RA3-6B2, 12-0452-82, eBioscience), CD11b (BV510, clone M1/70, 101263, Biolegend), CD11c (BV711, clone N418, 117349, Biolegend), F4/80 (APC, clone BM8, 123116, Biolegend), CD80 (PE-Cy7, clone 16-10A1, 104734, Biolegend) and CD86 (BV605, clone GL1, 105037, Biolegend).

### Immunohistology staining of the lymph node

The draining lymph nodes were snap frozen with liquid nitrogen by embedding the tissue into the OCT compound in a Cryomold (Tissue-Tek) and then cryosectioned. The tissue slices (10 µm thickness) were stained with CD3e (1:50, APC, clone 145-2C11, 17-0031-82 eBioscience), B220 (1:50, PE, clone RA3-6B2, 14-0452-82, eBioscience) and Hoechst 33342 dye (H1399, Thermofisher). The whole lymph node was scanned using the Leica SP8 Spectral Confocal Microscope. The images from individual channels were processed using Fiji software.

### In vivo immunization studies

For the SIINKFEL vaccination studies, two doses of RP phages displaying the SIINFEKL pVIIIs (5*×*10^12^ phage particles calculated based on a ssDNA length of 7234 nucleotides, 100 μL 1*×* PBS), or an equivalent amount of free SIINKFEKL peptide ( ∼ 3.8 nmol, ∼ 3.7 μg) mixed with CpG oligonucleotide ( ∼ 3.6 nmol, 5’-TCCATGACGTTCCTGACGTT-3’, Integrated DNA Technologies) were administrated via tail base injection on day 0 and day 7. On day 14, ∼ 30 μL of blood was used to study the antigen-specific CD8^+^ T cell response. Specifically, red blood cells (RBCs) were lysed using 300 μL of RBC lysis buffer (R7757, Millipore Sigma). The peripheral blood monocytes (PBMCs) were then collected by centrifugation at 500 g for 5 minutes and the supernatant was discarded. An additional 300 µL of RBC lysis buffer was added to further lyse any remaining RBCs, followed by a second centrifugation at 500 g for 5 minutes. After discarding the supernatant, the PBMCs were redispersed in 100 μL 1× PBS containing LIVE/DEAD™ fixable near-IR dye (L34976, Themofisher) for live/dead staining at room temperature for 10 minutes. The cells were centrifuged at 500 g for 5 minutes and then redispersed in 100 μL FACS buffer (1% BSA in 1× PBS) containing Fc blocker (101320, Biolegend), SIINFEKL H-2K^b^ tetramer (PE, TB-5001-1, MBL international) and anti-mouse CD8*α*antibody (APC, GTX76346, GENETEX) at room temperature for 30 minutes. Excess antibodies were removed through two centrifugation steps at 500 g for 5 minutes each.

For the Adpgk neoantigen in vivo vaccination study, RP phages displaying the Adpgk pVIIIs (5*×*10^12^ phage particles, calculated based on a ssDNA length of nucleotides, 100 μL 1*×* PBS), or equivalent amount of free Adpgk peptide (∼ 15.8 μg, Elim Biopharmaceuticals) mixed with CpG oligonucleotide (∼ 3.6 nmol) were administrated via tailbase injection. The H-2D^b^ Adpgk neoepitope Tetramer (PE, TB-5113-1, MBL international) and the anti-mouse CD279 (PD-1) Antibody (BV421, clone 29F1A12, 135221, Biolegend) were used for the cell staining.

For prophylactic tumor challenge studies, on day 15, B16F10-OVA cells or MC-38 cells in 100 μL 1× PBS were subcutaneously inoculated at the left flank of the mice. Tumor size was monitored every 2-4 days and calculated using the empirical formula: 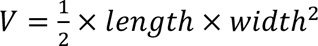. Mice were euthanized if the tumor volume exceeded 1000 mm^3^ or when animals became moribund with severe weight loss or ulceration.

### Cancer immunotherapy studies

For the MC-38 colon cancer treatment study, MC-38 colon cancer was first established by subcutaneous injection of 0.25 million cells (in 100 μL 1*×* PBS) in the left flank of the mouse. 8 days post tumor inoculation when the average tumor size reached ∼ 80 mm^3^, the mice were regrouped to make sure each study group had similar size distributions. RP phages (1.0*×*10^13^ phage particles, calculated based on a ssDNA length of 7234 nucleotides, 100 μL 1*×*PBS) or equivalent combinations of free Adpgk peptide (∼ 31.6 μg) and CpG (∼ 7.2 nmol) were administered in two doses on day 8 and 15 post tumor inoculation. Anti-mouse PD-1 antibodies (100 μg, clone RMP1-14, BioXcell, 1*×* PBS) were administered on day 2 and day 4 following each vaccination treatment, or alone at the same time. Tumor size was monitored every 2-4 days and calculated using the empirical formula: 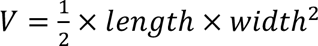. Mouse were euthanized if the tumor volume exceeded 1000 mm^3^ or when animals became moribund with severe weight loss or ulceration.

## Data availability

All data supporting the findings of this study are provided within the Article and Supplementary Information. Additional data are available from the corresponding author upon reasonable request.

