## supplementary information for "Reprogramming the genome of M13 bacteriophage for all-in-one personalized cancer vaccine"

### Supplementary Table 1: Reprogrammed ssDNA sequence

| Name | Sequence (5'→3') |
|---|---|
| CG0 | aatagtggactcttgttccaaactggaacaacactcaaccctatctcgggctattcttttgatttataagggattttgccgatttcggc<br>ctattggttaaaaaatgagctgatttaacaaaaatttaacgcgacatgtTGATATGCCTAAGGCAAGTTGCCAT<br>GTGCTGTGGGCTATCAGGCAGATTAGCTCTGATGCTCTAGCAGAGGCCATAGCAATA<br>GGCATTTGCTGGAGGCATCAGCCAAGGCCAGGTGCCAAAGGCCCCTTGCCAGGAGC<br>CAGTGGCCTTACAGCAATGAGCATCTTTGCTAGCTGCCTTTGTGCCATAGGCTAGGG<br>CATGGAGCTACAGGCAAGAGAGCTTTCTAGCTCCCAGCCAATTTGCTTCAAGCACCT<br>GCCATCAAGCTATTGCAAACAGGCCTTTGGGCTTTCCAGCACACTGCAGACTTGCCTT<br>GAGCCTACATGCATGCTGCCCCAGCACTGTGCCAGAAGCAGGTTTGCATGTCAGCTT<br>AGTGCATATCAGCTTTGGCCATGAGCACCTTGCTCTGTAGCAGAAGAGCATCTAGGC<br>ATTTGGCAATAGTGCTCTGAGCCAAATGCTGCTATGCTGCACTGCCTGTGGCAACAG<br>GCTTAAGAGCCCTTGAGCTTGGCAGCACCAGTGCTTTTGGCAATTGCAACTAGGCCA<br>AGAGGCAAGTCAGCCAGCCAGCAGATCAGCTAGCTAGCAGTAGGCATGTGCTACTGT<br>GCAAAATGGCTAAAAAGCCACTGGGCCAGGGGCTTAACTGCACAAGGGCTAAGGAG<br>CTAAAAGCTCTATGCTCATGCAGAGAAGCTATGCTGCTCTGGGCTTGTTGCCCTCTGC<br>ACCTGGGCTTGGGGCTTGATGCATTAGGCTCTAGGCTGAAGGCTCAGAGCTGGACAG<br>CCTCCTGGCATTAAGCTTGAGTGCATAGGGCACCTGGCTAACTAGCATGGTGCAGGA<br>GGCCTAGTAGCATTGGCTAATAGGCAGGATGCTTATGCATATAGCATGGTGGCTGTAT<br>GCATGTGCAGACAGCATTAGTGCTCAGGAGCAAAGGGCTACCTGCCCAATGGCCAAC<br>TGCTTGTAGCCACCAAGCATTCAGGCgcggccgcacgcgatacaatccgcgccctgtagcggcgcattaa<br>gcgcggcgggtgtggtggttacgcgcagcgtgaccgctacacttgccagcgccctagcgcccgctcctttcgctttcttcccttcc<br>tttctcgccacgttcgccggctttccccgtcaagctctaaatcgggggctccctttagggttccgatttagtgctttacggcacctcg<br>accccaaaaaacttgatttgggtgatggttcacgtagtgggccatcgccctgatagacggtttttcgccctttgacgttggagtcca<br>cgttcttt |
| CG09 | aatagtggactcttgttccaaactggaacaacactcaaccctatctcgggctattcttttgatttataagggattttgccgatttcggc<br>ctattggttaaaaaatgagctgatttaacaaaaatttaacgcgacatgtTGATATCGCTAAGCGAAGTTGCCAT<br>GTGCTGTGGGCTATCAGGCAGATTAGCTCTGATCGTCTAGCAGAGCGCATACGAATA<br>GGCATTTGCTGGAGGCATCAGCCAAGGCCAGGTGCCAAAGGCCCCTTGCCAGGAGC<br>CAGTGCGCTTACACGAATGAGCATCTTTGCTAGCTGCCTTTGTGCCATAGCGTAGGC<br>GATGGAGCTACAGGCAAGAGACGTTTCTAGCTCCCACGCAATTTCGTTCAAGCACCT<br>CGCATCAACGTATTGCAAACAGCGCTTTGGGCTTTCCACGACACTCGAGACTTGCCTT<br>GACGCTACATCGATGCTGCCCCAGCACTGTCGCAGAACGAGGTTTGCATGTCAGCTT<br>AGTCGATATCAGCTTTGGCCATGAGCACCTTCGTCTGTAGCAGAAGAGCATCTAGGC<br>ATTTGGCAATAGTGCTCTGAGCCAAATGCTGCTATGCTGCACTGCCTGTGCGAACAG<br>GCTTAAGAGCCCTTGAGCTTGGCACGACCAGTGCTTTTGGCAATTGCAACTAGCGCA<br>AGAGCGAAGTCAGCCAGCCACGAGATCACGTAGCTAGCAGTAGGCATGTGCTACTGT<br>GCAAAATGGCTAAAAAGCCACTGGGCCAGGGCGTTAACTCGACAAGGCGTAAGGAC<br>GTAAAAGCTCTATGCTCATGCAGAGAAGCTATGCTCGTCTGGGCTTGTTGCCCTCTGC<br>ACCTGGCGTTGGGCGTTGATGCATTAGCGTCTAGCGTGAAGGCTCAGAGCTGGACAC<br>GCTCCTGGCATTAAGCTTGAGTCGATAGGGCACCTGGCTAACTACGATGGTGCAGGA<br>GGCCTAGTAGCATTGCGTAATAGGCAGGATGCTTATGCATATAGCATGGTGGCTGTAT<br>GCATGTGCAGACAGCATTAGTGCTCAGGACGAAAGGGCTACCTCGCCAATGGCCAAC<br>TCGTTGTAGCCACCAAGCATTCAGGCgcggccgcacgcgatacaatccgcgccctgtagcggcgcattaa<br>gcgcggcgggtgtggtggttacgcgcagcgtgaccgctacacttgccagcgccctagcgcccgctcctttcgctttcttcccttcc<br>tttctcgccacgttcgccggctttccccgtcaagctctaaatcgggggctccctttagggttccgatttagtgctttacggcacctcg<br>accccaaaaaacttgatttgggtgatggttcacgtagtgggccatcgccctgatagacggtttttcgccctttgacgttggagtcca<br>cgttcttt |
| CG27 | aatagtggactcttgttccaaactggaacaacactcaaccctatctcgggctattcttttgatttataagggattttgccgatttcggc<br>ctattggttaaaaaatgagctgatttaacaaaaatttaacgcgacatgtTGATATCGCTAAGCGAAGTTCGCAT<br>GTCGTGTGGCGTATCAGCGAGATTACGTCTGATCGTCTACGAGAGCGCATACGAATA<br>GCGATTTCGTGGAGCGATCACGCAAGCGCAGGTCGCAAAGCGCCCTTCGCAGGACG<br>CAGTGCGCTTACACGAATGACGATCTTTCGTAGCTCGCTTTGTCGCATAGCGTAGGC<br>GATGGACGTACAGCGAAGAGACGTTTCTACGTCCCACGCAATTTCGTTCAACGACCT<br>CGCATCAACGTATTCGAAACAGCGCTTTGGCGTTTCCACGACACTCGAGACTTCGCTT |

| | |
|---|---|
| | GACGCTACATCGATGCTCGCCCACGACTGTCGCAGAACGAGGTTTCGATGTCACGTT<br>AGTCGATATCACGTTTGCGCATGACGACCTTCGTCTGTACGAGAAGACGATCTAGCG<br>ATTTGCGAATAGTCGTCTGACGCAAATCGTGCTATCGTGCACTCGCTGTGCGAACAG<br>CGTTAAGACGCCTTGACGTTGGCACGACCAGTCGTTTTGCGAATTCGAACTAGCGCA<br>AGAGCGAAGTCACGCAGCCACGAGATCACGTAGCTACGAGTAGCGATGTCGTACTGT<br>CGAAAATGCGTAAAAACGCACTGGCGCAGGGCGTTAACTCGACAAGGCGTAAGGAC<br>GTAAAACGTCTATCGTCATCGAGAGAACGTATGCTCGTCTGGCGTTGTTCGCCTCTCG<br>ACCTGGCGTTGGGCGTTGATCGATTAGCGTCTAGCGTGAAGCGTCAGACGTGGACAC<br>GCTCCTGCGATTAACGTTGAGTCGATAGGCGACCTGCGTAACTACGATGGTCGAGGA<br>GCGCTAGTACGATTGCGTAATAGCGAGGATCGTTATCGATATACGATGGTGCGTGTAT<br>CGATGTCGAGACACGATTAGTCGTCAGGACGAAAGGCGTACCTCGCCAATGCGCAAC<br>TCGTTGTACGCACCAACGATTCAGCGgcggccgcacgcgatacaatccgcgccctgtagcggcgcattaa<br>gcgcggcgggtgtggtggttacgcgcagcgtgaccgctacacttgccagcgccctagcgcccgctcctttcgctttcttcccttcc<br>tttctcgccacgttcgccggctttccccgtcaagctctaaatcgggggctccctttagggttccgatttagtgctttacggcacctcg<br>accccaaaaaacttgatttgggtgatggttcacgtagtgggccatcgccctgatagacggtttttcgccctttgacgttggagtcca<br>cgttcttt |
| CpG40 | aatagtggactcttgttccaaactggaacaacactcaaccctatctcgggctattcttttgatttataagggattttgccgatttcggc<br>ctattggttaaaaaatgagctgatttaacaaaaatttaacgcgacatgtTGATATCGCTAAGGACGTTAAGTTC<br>GCATGTGACGTTTGTGGCGTATCAGCGAGATTAGACGTTTCTGATAACGTTTCTACGA<br>GAGGACGTTCATACGAATAGGACGTTATTTCGTGGAGCGATCAGACGTTCAAGAACG<br>TTCAGGTCGCAAAGCGCCCTTGACGTTCAGGAAACGTTCAGTGCGCTTACACGAATG<br>ACGATCTTTCGTAGCTCGCTTTGTAACGTTCATAGCGTAGGGACGTTATGGACGTACA<br>GAACGTTAAGAGACGTTTCTAGACGTTTCCCAGACGTTCAATTTCGTTCAACGACCTC<br>GCATCAAAACGTTTATTCGAAACAGCGCTTTGGCGTTTCCACGACACTCGAGACTTCG<br>CTTGAGACGTTCTACATCGATGCTAACGTTCCCAGACGTTACTGTGACGTTCAGAACG<br>AGGTTTCGATGTCACGTTAGTCGATATCAGACGTTTTTGGACGTTCATGACGACCTTA<br>ACGTTTCTGTACGAGAAGACGATCTAGCGATTTGGACGTTAATAGTCGTCTGAGACGT<br>TCAAATCGTGCTATCGTGCACTGACGTTCTGTGGACGTTAACAGCGTTAAGACGCCTT<br>GACGTTGGCACGACCAGTCGTTTTGGACGTTAATTCGAACTAGCGCAAGAGCGAAGT<br>CACGCAGCCAAACGTTAGATCACGTAGCTAGACGTTAGTAGGACGTTATGTCGTACTG<br>TCGAAAATGCGTAAAAACGCACTGGCGCAGGGCGTTAACTGACGTTACAAGGCGTAA<br>GGACGTAAAACGTCTATCGTCATCGAGAGAACGTATGCTGACGTTTCTGGCGTTGTTA<br>ACGTTCCTCTCGACCTGGCGTTGGGGACGTTTTGATCGATTAGCGTCTAGGACGTTT<br>GAAGCGTCAGAGACGTTTGGACACGCTCCTGCGATTAACGTTGAGTAACGTTATAGG<br>GACGTTACCTGCGTAACTAGACGTTATGGTCGAGGAGCGCTAGTAGACGTTATTGCG<br>TAATAGCGAGGATAACGTTTTATGACGTTATATAAACGTTATGGTGCGTGTATCGATGT<br>CGAGACAAACGTTATTAGTCGTCAGGAGACGTTAAAGGGACGTTTACCTGACGTTCCA<br>ATGAACGTTCAACTCGTTGTAGACGTTCACCAACGATTCAGCGgcggccgcacgcgatacaat<br>ccgcgccctgtagcggcgcattaagcgcggcgggtgtggtggttacgcgcagcgtgaccgctacacttgccagcgccctagc<br>gcccgctcctttcgctttcttcccttcctttctcgccacgttcgccggctttccccgtcaagctctaaatcgggggctccctttagggttc<br>cgatttagtgctttacggcacctcgaccccaaaaaacttgatttgggtgatggttcacgtagtgggccatcgccctgatagacggt<br>ttttcgccctttgacgttggagtccacgttcttt |

*Gblock is highlighted in capital letters.

### Supplementary Table 2: pVIII sequence of the helper plasmid

#### Supplementary Table 2.1.: Full amino acid sequence of the pVIII

| helper plasmid | location | pVIII type | mature pVIII sequence |
|---|---|---|---|
| rEES | recombinant pVIII | antigen pVIII | AEESIINFEKL#DPAKAAFDSLQASATEYIGYAW<br>AMVVVIVGATIGIKLFKKFTSKAS |
| | original pVIII | wild-type pVIII | AEGDDPAKAAFDSLQASATEYIGYAWAMVVVIV<br>GATIGIKLFKKFTSKAS |
| EES | original pVIII | antigen pVIII | AEESIINFEKLDPAKAAFDSLQASATEYIGYAWAM<br>VVVIVGATIGIKLFKKFTSKAS |
| | recombinant pVIII | wild-type pVIII | AEGDDPAKAAFDSLQASATEYIGYAWAMVVVIV<br>GATIGIKLFKKFTSKAS |
| EESmut | original pVIII | antigen pVIII | AEESIINFEKLDPAKAAFDSLQASATEYIGYAWA<br>MVVVIVGATIGIKLFKKFTSKAS |
| | recombinant pVIII | wild-type pVIII | AEGDDPAKAAFDSLQASATEYIGYAWAMVVVIV<br>GATIGIKLFKKFTSKAS |
| rAE | recombinant pVIII | antigen pVIII | ASMTNMELMEDPAKAAFDSLQASATEYIGYAW<br>AMVVVIVGATIGIKLFKKFTSKAS |
| | original pVIII | wild-type pVIII | AEGDDPAKAAFDSLQASATEYIGYAWAMVVVIV<br>GATIGIKLFKKFTSKAS |
| AE | original pVIII | antigen pVIII | ASMTNMELMEDPAKAAFDSLQASATEYIGYAW<br>AMVVVIVGATIGIKLFKKFTSKAS |
| | recombinant pVIII | wild-type pVIII | AEGDDPAKAAFDSLQASATEYIGYAWAMVVVIV<br>GATIGIKLFKKFTSKAS |

#: antigen sequences underlined and highlighted in purple.

#### Supplementary Table 2.2.: Nucleotide sequence the selected pVIII (including the 20 nucleotides proceeding the start codon of the pVIII)

| helper plasmids | location | pVIII type | Nucleotide sequence of the pVIII (5'⟶3', including 20 bases proceeding the start codon) |
|---|---|---|---|
| EES | original pVIII | antigen pVIII | CGTTTAATGGAAGCTTCCTCATG^AAAAAGTCTTTAG<br>TCCTCAAAGCCTCTGTAGCCGTTGCTACCCTCGTTC<br>CGATGCTGTCTTTCGCTGCTGAGGAAAGTATAATCA<br>ACTTTGAAAAACTGGATCCCGCAAAAGCGGCCTTTG<br>ACTCCCTGCAAGCCTCAGCGACCGAATATATCGGTT<br>ATGCGTGGGCGATGGTTGTTGTCATTGTCGGCGCA<br>ACTATCGGTATCAAGCTGTTTAAGAAATTCACCTCG<br>AAAGCAAGCTGA |
| EESmut | original pVIII | antigen pVIII | CGTTTAATGGT*AGCTTCCTCATG^AAAAAGTCTTTA<br>GTCCTCAAAGCCTCTGTAGCCGTTGCTACCCTCGTT<br>CCGATGCTGTCTTTCGCTGCTGAGGAAAGTATAATC<br>AACTTTGAAAAACTGGATCCCGCAAAAGCGGCCTTT<br>GACTCCCTGCAAGCCTCAGCGACCGAATATATCGG<br>TTATGCGTGGGCGATGGTTGTTGTCATTGTCGGCG<br>CAACTATCGGTATCAAGCTGTTTAAGAAATTCACCT<br>CGAAAGCAAGCTGA |

*mutation underlined and highlighted in grey. ^start codon of pVIII underlined.

### Supplementary Figure 1. SIINFEKL pVIII characterization with HPLC and MALDI-TOF mass spectrometry

SIINFEKL pVIII-expressing RP phages were amplified with an RP phage phagemid and the rEES helper plasmid. Various fractions (fxns) were collected in Supplementary Fig. 1a at the characteristic peaks as indicated by different colors and analyzed with MALDI - TOF MS. The main peak at ~ 55.8 min (fxn 56 and part of fxn 57) was identified to be the wild-type pVIII with molecular weight of 5329 Da and the shoulder peak following it at ~ 56.5min (part of fxn 57) was the SIINFEKL pVIII with molecular weight of 6142 Da.

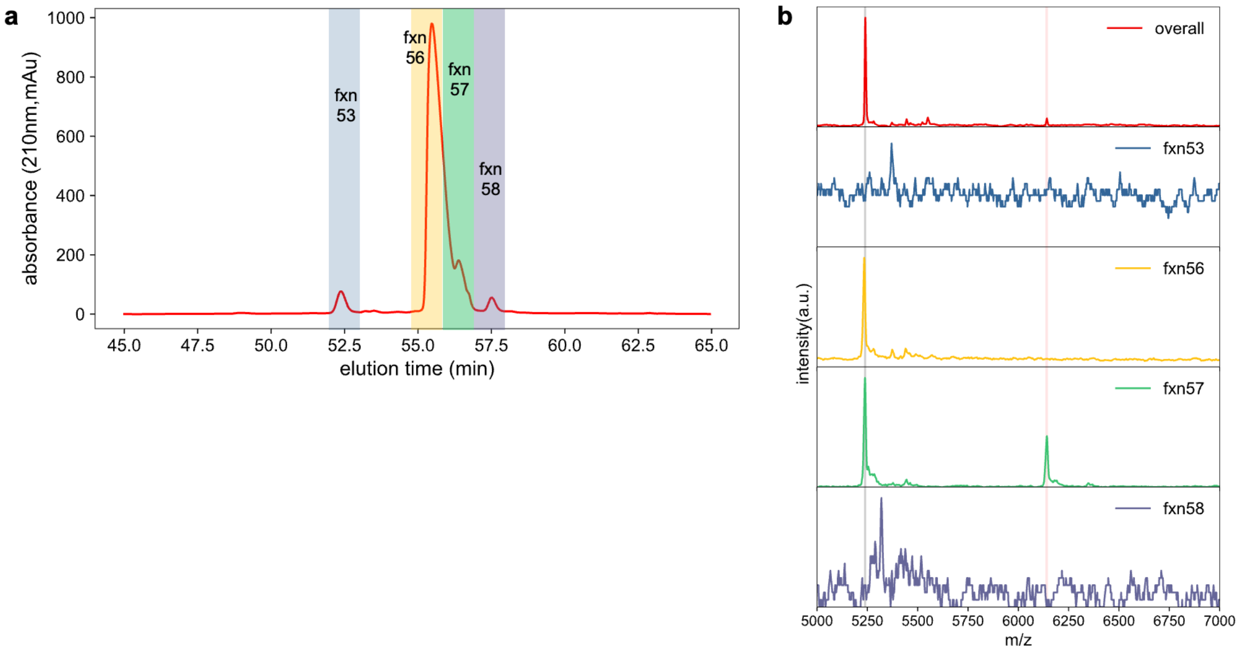


**Supplementary Fig. 1 | Identification of SIINFEKL pVIII with HPLC and MALDI-TOF MS. a**, Various factions of the SIINFEKL pVIII-displaying phages were collected with HPLC as indicated by different colors. **b**, MALDI-TOF MS plot of the various HPLC fractions collected in (**a**).

**Supplementary Figure 2. FAM conjugation and quantification on the RP phages**

The fluorescein (FAM) density on the phage surface (number of FAM molecules per phage particle) was quantified by using absorption measurement. First, a linear calibration curve of absorbance vs. dye concentration was constructed by measuring the absorbance of the free FAM at various concentrations as in Supplementary Figs 2a,b. To characterize the FAM density on the phage surface, the absorption spectrum of the phage-FAM was measured making sure that the peak absorbance falls in the linear absorption regime (Supplementary Fig. 2c as the normalized absorption spectrum of the phage-FAM complex). Then the FAM concentration was determined from the calibration curve and the FAM density on the phage surface was determined as $[FAM](M)/[phage](M)$. For a typical phage-FAM complexation study, on average there were ~ 680 FAM molecules conjugated on each phage (the phage particle concentration was calculated based on the ssDNA length of 7234 nucleotides).

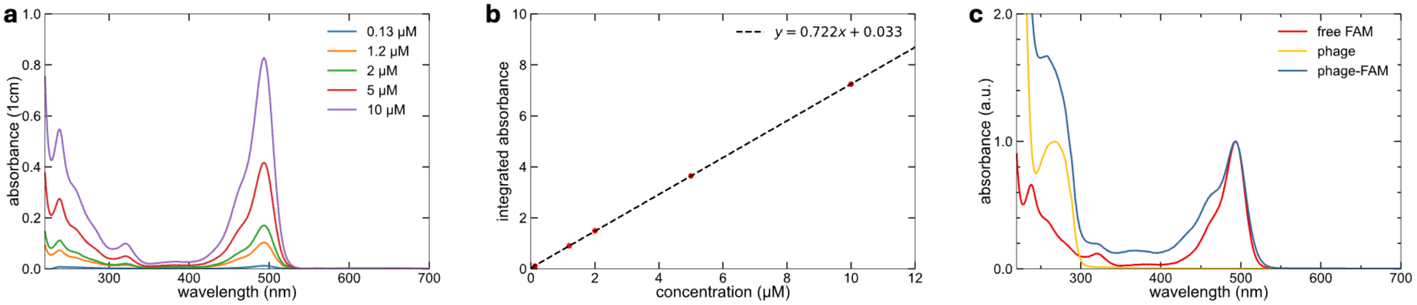


**Supplementary Fig. 2 | Quantification of the FAM density on the RP phage with absorption measurement.** **a**, Absorption spectrum of free FAM in 1× PBS at various concentrations. **b**, Linear calibration curve of the absorbance vs. concentration for the free FAM. **c**, Normalized absorption spectrum of phage, free FAM and phage-FAM complex confirmed the conjugation of FAM on the phage surface.

**Supplementary Figure 3. FACS gating of the phage distribution among immune cells and activation of immune cells in draining lymph nodes**

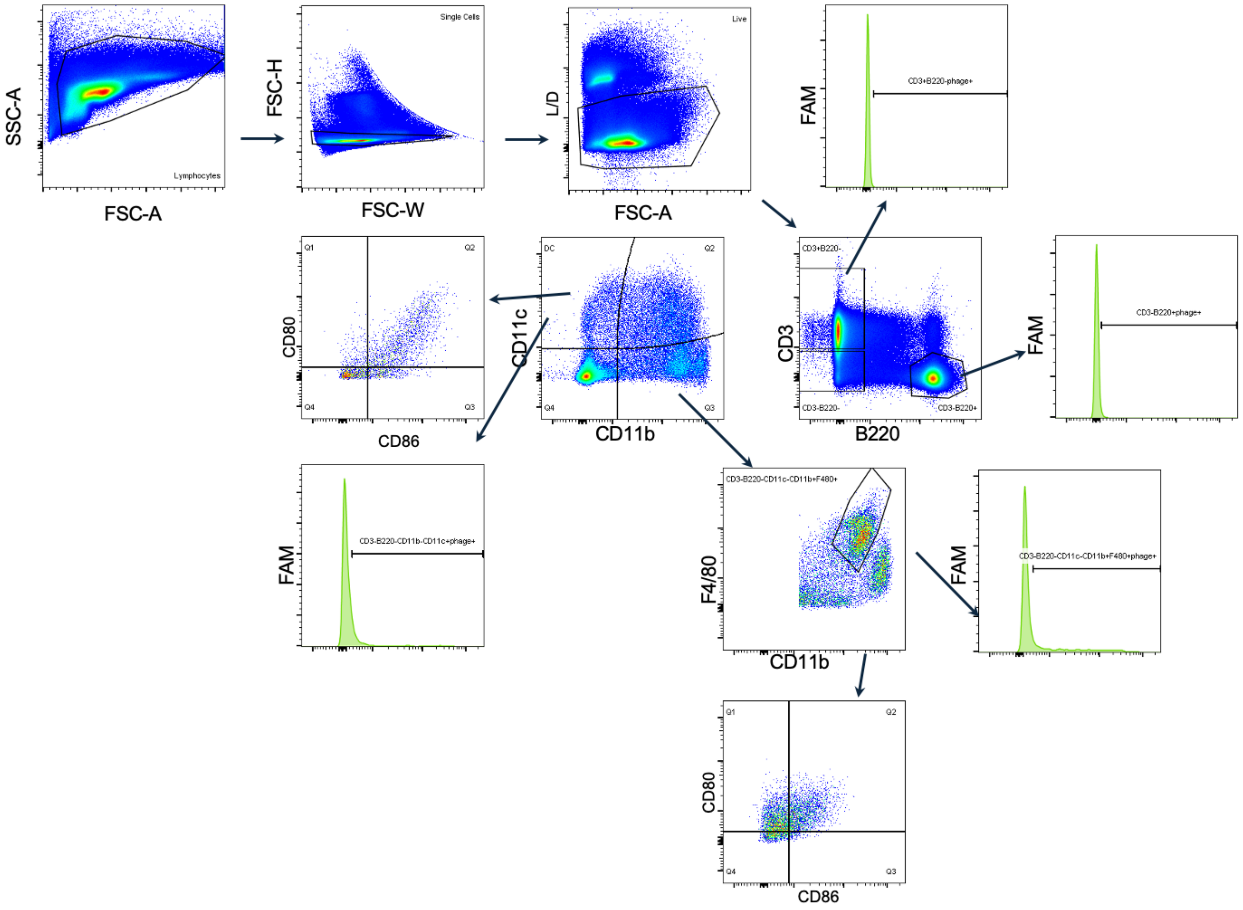

### Supplementary Figure 4. FACS gating of the antigen-specific CD8$^+$ memory T cells

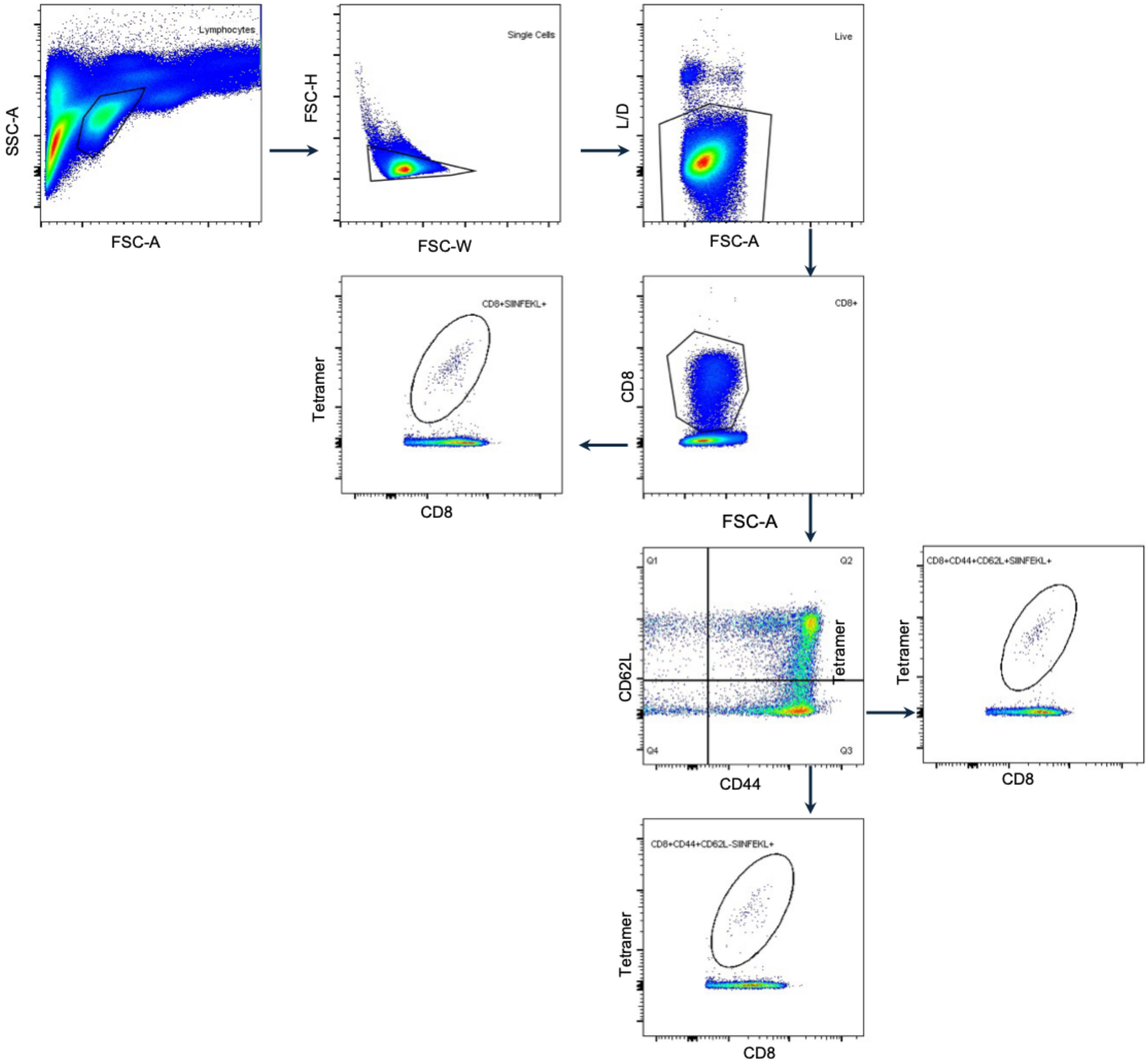

**Supplementary Figure 5. cDNA sequencing of the mouse Adpgk gene**

The total RNA of MC-38 cells was extracted using the RNeasy Mini Kit (74104, Qiagen). The first-strand cDNA was synthesized using 5$\mu$g of the total RNA with the SuperScript III First-Strand Synthesis SuperMix (18080400, Invitrogen). Adpgk cDNA with lengths of 555 bp were amplified using the forward primer:5'-CTGGAGGTGTTTGTGTCTAG-3' and reverse primer 5'-TCTTAGAGACACTCGGTTGG-3' with the Q5® High-Fidelity 2× Master Mix (New England Biolabs). The length of the PCR product was confirmed with 2% agarose gel and the sequence was confirmed with Sanger sequencing using the forward primer as the sequencing primer.

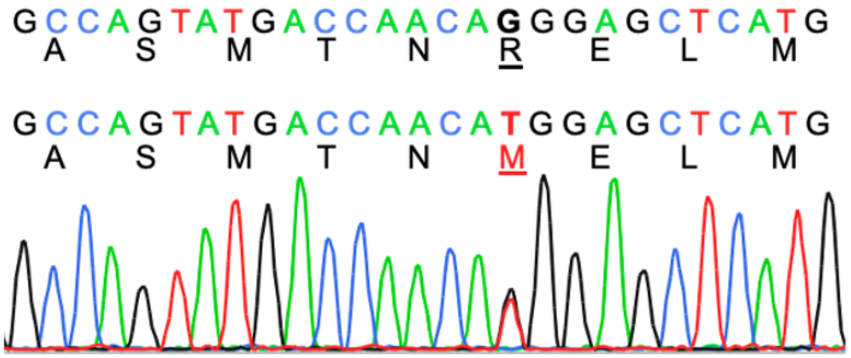

### Supplementary Figure 6. HPLC collection and MALDI-TOF MS analysis of the Adpgk RP phage

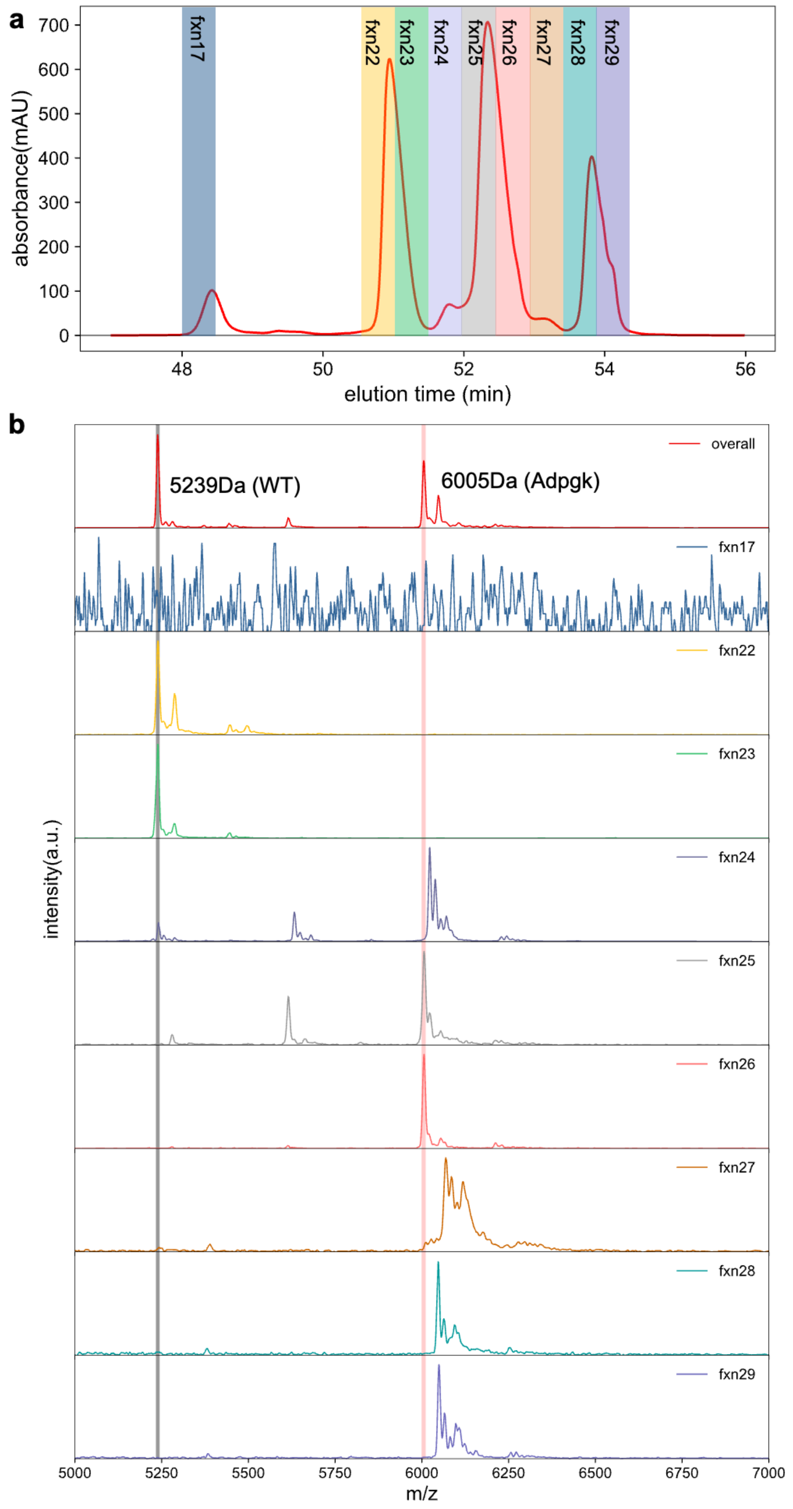


**Supplementary Fig. 6 | Identifying the Adpgk pVIII peak with HPLC and MALDI-TOF MS. a**, HPLC collection plot of the Adpgk phage with the shading areas indicating the fractions that were collected for MALDI-TOF analysis. **b**, MALDI-TOF analysis of the collected fractions in (**a**). The results show that fractions 22,23 and 25,26 were wild-type pVIII (5239Da) and the Adpgk pVIII (6005Da), respectively.

### Supplementary Figure 7. Low neoantigen-specific CD8⁺ T cell response of the Adpgk RP phages due to low adjuvanticity and antigen display ratio

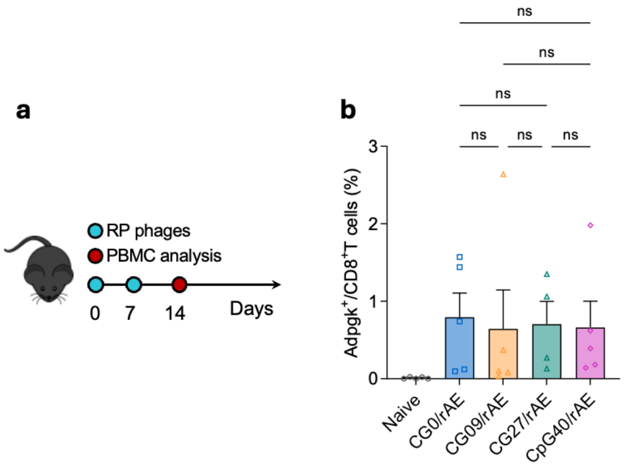


**Supplementary Fig. 7 | Initial trial with Adpgk pVIII-displaying RP phages showed low immune response. a**, Dosing schedule of the initial vaccination study with the RP phages amplified by combining different RP phagemids and the rAE helper plasmid. **b**, The resultant low neoantigen-specific CD8$^{+}$ T response as studied by the tetramer staining. ns: none significance, analyzed by two-way ANOVA (**b**,**e**) with Bonferroni post hoc test.

### Supplementary Figure 8. Treatment timing is critical for combination therapy outcomes

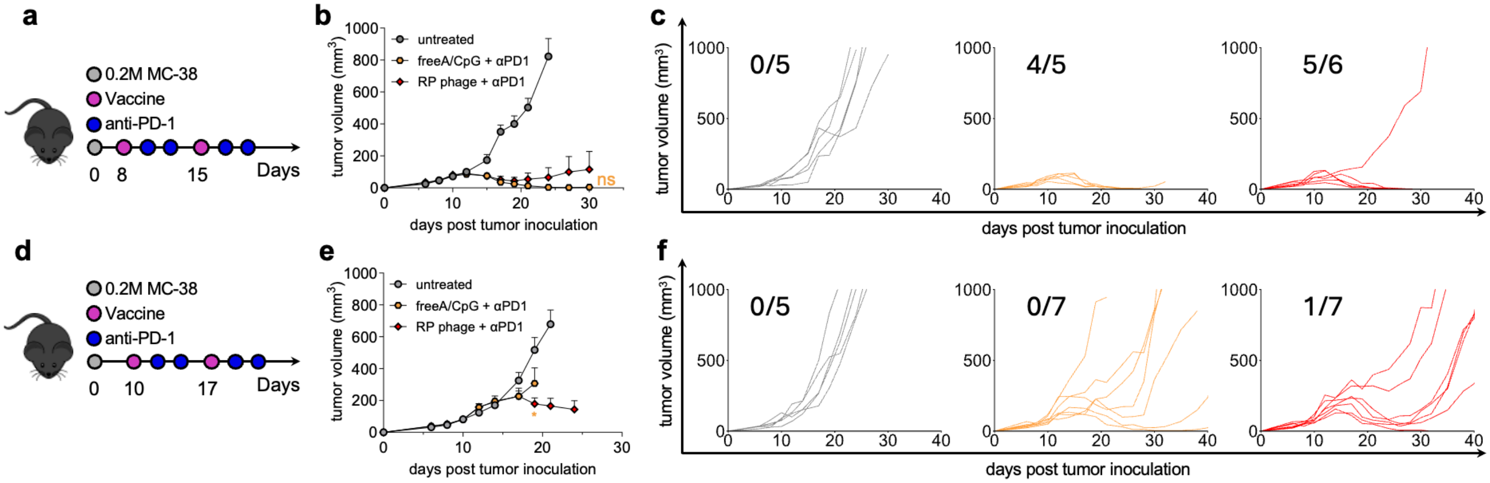


**Supplementary Fig. 8 | Combination therapy efficacy depended strongly on the treatment timing.** **a**,**b**,**c**, Combination treatment started at day 8 yielded high complete regression rate, with 0.2 M MC-38 inoculation when the average tumor volume was ~ 45 $mm^3$. **a**, RP phages ($7.5\times10^{12}$ phage particles calculated based on a ssDNA length of 7234 nucleotides, 100 μL 1× PBS) or equivalent combinations of free Adpgk peptide (~ 23.7 μg) and CpG (~ 5.4 nmol) were administered in two doses on day 8 and 15 post tumor inoculation. Anti-mouse PD-1 antibodies (100 μg) were administered on day 2 and 4 following each vaccination treatment. **b**, Average tumor volume of the combined therapy study and (**c**) individual MC-38 tumor growth curves. **d**,**e**,**f**, Combination treatment started at day 9 yielded low complete regression rate, with 0.2M MC-38 inoculation when the average tumor volume was ~ 80 $mm^3$. **d**, RP phages ($7.5\times10^{12}$ phage particles calculated based on a ssDNA length of 7234 nucleotides, 100 μL 1× PBS) or equivalent combinations of free Adpgk peptide (~ 23.7 μg) and CpG (~ 5.4 nmol) were administered in two doses on day 10 and 17 post tumor inoculation. Anti-mouse PD-1 antibodies (100 μg) were administered on day 2 and 4 following each vaccination treatment. **e**, Average tumor volume of the combined therapy study and (**f**) individual MC-38 tumor growth curves. ns: none significance, $*p<0.05$, analyzed by two-way ANOVA (**b**,**e**) with Bonferroni post hoc test.

In the course of evaluating combination therapy, we identified the complicated dependency of therapeutic efficacy on the timing of treatment administration. Specifically, for the same tumor inoculation dose, early-stage treatment starting at day 8, when the average tumor size was ~ 45 $mm^3$, yielded a complete regression rate exceeding 80%. In contrast, administration at a later stage of day 10, when the average tumor size was ~ 80 $mm^3$, substantially diminished therapeutic outcomes (Supplementary Fig. 8). These findings emphasize the importance of early tumor diagnosis and highlight the challenges associated with managing advanced malignancies.
